# Systematic and proactive evaluation of AIRE missense variant effects

**DOI:** 10.1101/2025.11.18.688830

**Authors:** Anna Axakova, Amund H. Berger, Warren van Loggerenberg, Nishka Kishore, Marinella Gebbia, Megan X. Ding, Samuel V. Douville, Daniel R. Tabet, Atina G. Cote, Jochen Weile, Stefan Johansson, Eirik Bratland, Frederick P. Roth

**Affiliations:** Donnelly Centre for Cellular and Biomolecular Research, University of Toronto, Toronto, Ontario, Canada; Department of Molecular Genetics, University of Toronto, Toronto, Ontario, Canada; Lunenfeld-Tanenbaum Research Institute, Sinai Health, Toronto, Ontario, Canada; Department of Clinical Science, University of Bergen, Norway; Department of Computational and Systems Biology, University of Pittsburgh School of Medicine, Pittsburgh, Pennsylvania, USA; Faculty of Health Science, McMaster University, Hamilton, Ontario, Canada; Department of Medical Genetics, Haukeland University Hospital, Bergen, Norway

## Abstract

Pathogenic variants in the Autoimmune Regulator (*AIRE*) gene cause Autoimmune Polyendocrine Syndrome Type 1 (APS-1), a rare primary immunodeficiency disease with symptoms including hypoparathyroidism, adrenal insufficiency, and chronic mucocutaneous candidiasis. AIRE increases the expression and presentation of tissue-specific genes expressing ‘self’ antigens in the developing T cell niche, thus triggering the elimination of self-reactive T cells and preventing autoimmunity. Earlier diagnoses can benefit patients, and APS-1 diagnosis by AIRE sequencing is increasingly common. However, two thirds of reported clinical variants are missense, and more than half of these are “variants of uncertain significance” (VUS). Cell-based variant functional assays can provide strong evidence towards more informative variant classification, but these are carried out reactively, often years after clinical presentation. By contrast, proactively assessing all possible missense variants could provide immediate evidence to guide genetic diagnosis, even for never-before-seen variants. Here we used an insulin promoter-driven reporter to proactively assess the function of 9790 *AIRE* missense variants. The resulting *AIRE* variant effect map both validates and extends current biochemical knowledge, concords with pathogenicity annotations, and provides proactive evidence for 70% of previously-reported VUS. Placing our map in the context of both an international APS-1 cohort and the UK BioBank revealed quantitative genotype-phenotype correlations. Using current guidelines, we provide classifications for 32% of current VUS. Together, our proactive resource of AIRE variant impacts offers the potential to improve patient outcomes via more rapid and definitive APS-1 diagnosis.

## Introduction

Autoimmune Polyendocrine Syndrome Type 1 (APS-1, MIM:240300), also called Autoimmune Polyendocrinopathy with Candidiasis and Ectodermal Dystrophy (APECED), is a rare primary immunodeficiency disease caused by variants in the autoimmune regulator (*AIRE*) gene^1^. Usually, APS-1 diagnosis requires the presence of two symptoms amongst hypoparathyroidism, adrenal insufficiency and chronic mucocutaneous candidiasis (henceforth “candidiasis”) but patients often present a broad range of other autoimmune symptoms, some of which may be life threatening^2^. Patients with APS-1 can have normal life expectancies with proper management of symptoms, but the prevalence of disease components increases over the years making treatment and follow-up problematic. Hence, earlier diagnosis improves patient outcomes by mitigating the effects of long-term autoimmunity^3^. Sequencing is increasingly used to diagnose APS-1 and proposes to decrease diagnostic delay^1,3^, but this in turn requires rapid and accurate interpretation of *AIRE* variants.

Of the *AIRE* variants reported to ClinVar^4^, a database of genetic variants and their impact on human health, two-thirds are missense variants, which are usually more challenging than other variants (e.g., nonsense, frameshift, or large deletion variants) to interpret. Indeed, 81% of *AIRE* missense variants in ClinVar are annotated as a “variant of uncertain significance”. One of the strongest forms of evidence for variant annotation under current guidelines, and one which is often unavailable, comes from functional assays of variant impact^5^. Currently, one-at-a-time ‘reactive’ variant functional assays are the standard, with results often lagging months or years after a rare variant has first been detected in a patient^6^. A more recent ‘proactive’ approach, made possible by advances in cellular engineering and sequencing, is to obtain functional evidence for nearly all possible missense variants (variant effect maps) that even include variants not yet seen in the clinic^6,7^.

Here we describe the first variant effect map for *AIRE*, developed using an insulin-driven GFP reporter in HEK293 cells. The map recapitulates known regions important for function (except partially the histone interacting PHD1 domain), provides evidence to guide clinical variant interpretation for APS-1, supports a genotype-phenotype correlation in an international patient cohort, associates apparently-damaging variants with hypoparathyroidism and vitamin B12 deficiency anemia in the UK BioBank, and allows for confident classification of 32% of VUS. The map reveals novel findings, e.g., many *AIRE* variants which show gain-of-function in terms of increased expression of target genes. The map also raises new questions, e.g., by identifying *AIRE* residues that are highly intolerant to substitution but for which the function remains unknown.

## Materials and Methods

### Plasmids

The Tet promoter of the dAAVS1-TetBxb1-BFP plasmid^8^ was replaced with the EF1α promoter from PL-sin-EF1α-eGFP (Addgene #21320) using Gibson Assembly^9^ to create the dAAVS1-EF1α-BFP plasmid. A Gateway-compatible attB integration construct (pDEST-Bxb1-25 Rec v2) was made by replacing the Kozak sequence and PTEN open reading frame of attB-PTEN-IRES-mCherry with the 1873bp Gateway attR1-CcdB-attR2 cassette from pDEST-AD-CYH2.

A codon-optimized version of the *AIRE* cDNA sequence was designed (see Supplemental Note for sequence; Twist Biosciences) because of the high GC-content within the endogenous *AIRE* sequence (NCBI Reference Sequence: NP_000374.1). To clone the *AIRE* DNA fragment into a plasmid backbone, the fragment was PCR-amplified with primers ATTB_U2site_kozak_F and ATTB_D1site_kozak_R (see Data S2 for Primers), with additional “universal addition” site needed for later variant library preparation, a Kozak site, and attB1 and attB2 sites on either end, compatible with integration into entry vector pDONR223. The PCR product was gel extracted, BP-cloned with Gateway™ BP Clonase™ II Enzyme mix (Thermo Fisher Scientific) into pDONR223 and transformed into NEB® 5-alpha Competent *E. coli* (High Efficiency) and selected on Spectinomycin LB plates. DNA was extracted with the QIAprep Spin Miniprep Kit and Sanger sequencing confirmed the plasmid. The AIRE WT plasmid compatible with integration into the *Bxb1* site was prepared by cloning AIRE WT in pDONR223 using Gateway™ LR Clonase™ into the *Bxb1*-compatible pDEST-HC-REC-v2, the plasmid enabling integration into the Bxb1 site of reporter cell lines (which also contains an downstream internal ribosomal entry site (IRES) driving expression of the mCherry integration marker). The product was transformed into 5-alpha *E.coli*, selected on LB plates with 1X Ampicillin, DNA extracted and sequence confirmed with Sanger.

To prepare for integrating the insulin promoter-driven GFP reporter construct into the Flp-In site of HEK293 cells, we adapted the pcDNA5-FRT-eGFP plasmid^10^ (kind gift from Anne-Claude Gingras) by replacing the existing CMV enhancer and promoter with the insulin core promoter. This was accomplished using PCR with phosphorylated primers delCMV_pcDNA5_FRT_eGFP_F and delCMV_pcDNA5_FRT_eGFP_R (see Data S2 for Primers) to linearize the plasmid. Next, the product was run on 1% agarose gel and band extracted with QIAQuick Gel Extraction Kit. The resulting product was ligated and transformed into NEB^®^ 5-alpha Competent (High Efficiency) *E.coli*, colonies picked and grown overnight at 37°C in LB with 1X Ampicillin selection, DNA extracted with QIAprep Spin Miniprep Kit, and Sanger sequenced to confirm. Next, to insert the insulin core promoter sequence, flanking Gateway-compatible attB sites were added to the core insulin sequence of -384 bp to 24 bp synthesized by Twist Bioscience (sequence available in Supplemental Note in Document S1) in a PCR reaction using primers INSpromATTB_F and INSpromATTB_R (see Data S2 for Primers). The resulting product was Gateway cloned into pDONR223 using Gateway™ BP Clonase™ II Enzyme mix, transformed, miniprepped, and subsequently LR gateway cloned using Gateway™ LR Clonase™ II Enzyme mix into ΔCMV-pcDNA5-FRT-eGFP to generate the final reporter construct plasmid InsProm-pcDNA5_FRT_eGFP. This final plasmid was transformed, DNA extracted and sequence confirmed with Sanger sequencing.

### Reporter cell line generation

To introduce the *Bxb1* landing pad for later introduction of *AIRE* variants, Flp-In™ T-REx™ HEK293 cells were transfected with equal proportions of dAAVS1-EF1α-BFP, AAVS1-TALEN-L and AAVS1-TALEN-R (Addgene #59025, #59026) using JetPRIME reagent (Polyplus Sartorius) as previously described^8,11^.

Transfected cells were cultured for ∼2 weeks and enriched for BFP-positive cells via fluorescence-activated cell sorting (FACS), cloned (by FACS), and validated as previously described^8,11^.

To insert the insulin reporter construct into the above Flp-In™ T-REx™ HEK293 EF1α-*Bxb1* cell line, 300K cells were seeded and co-transfected with 2 ug pOG44 (Flp-recombinase expression vector) and 200 ng InsProm-pcDNA5_FRT_eGFP using 4 uL JetPrime reagent and 200 uL of JetPrime buffer. To select for successful integrants, 200 ug/mL of Hygromycin B was added 48 hours after transfection. Surviving cells were seeded at 0.5 cells/well and clonally expanded. One clone was randomly selected as the insulin-driven GFP reporter cell line.

Cells were cultured in Serum (FBS), 90% Dulbecco’s Modified Eagle Medium (DMEM), and 1X Penicillin-Streptomycin (PS). Cells were tested for mycoplasma and grown at 37°C at 5% CO2.

### Small-scale reporter assay validation

As an initial test of the ability of the insulin-driven GFP reporter cell line to separate *AIRE* variants by function, we selected a small set of clinically-reported *AIRE* variants from across the length of the *AIRE* gene: p.Trp78Arg (pathogenic), p.Arg92Trp (pathogenic), p.Ser278Arg (benign), p.Pro400Leu (previously classified as benign, now with conflicting classifications), p.Ala403Thr (reported as both benign and VUS), and p.Asp503Asn (benign). Site-directed mutagenesis PCR was performed using the *AIRE* ORF in pDONR223 as a template to introduce for each variant (See Data S2 for list of site-directed mutagenesis primers, designed according to the codon-optimized AIRE sequence). Products were ligated, transformed into *E.coli*, and LR cloned into pDEST-HC-REC-v2 as described for the *AIRE* WT. Variants were confirmed using whole plasmid sequencing. To evaluate variants in the insulin-driven GFP reporter cell line, the reporter cell line was plated at 40K cells/well in DMEM + 10% FBS (antibiotic-free media) and variants transfected in triplicate using FuGENE^®^ 6 at a 10:1 ratio by mass of variant DNA:Bxb1 recombinase 30 minutes after plating the cells. Media was replaced with DMEM + 10% FBS + 1X PS 24 hours after transfection. Cells were expanded, gated for live, single cells which express the mCherry integration marker, and next evaluated for GFP expression one week after the transfection using a Beckman-Coulter flow cytometer to generate GFP distributions for the control variants.

### Library generation

We developed a modified version of the POPCode method^7^ called Dual-Tag Precision Oligo-Pool based Code Alteration (DT-POPCode). Relative to POPCode, DT-POPCode eliminates the need for a uracilated template and decreases variant bottlenecking during downstream plasmid library cloning (see Figure S1 and Supplemental Protocol in Document S1 for full protocol). First, mutagenic oligos were designed such that a “NNN” degenerate codon was centered on each *AIRE* target codon (codon-optimized, see Supplemental Note in Document S1 for sequence. Primers ATTB_U2site_kozak_F and ATTB_D1site_kozak_R were used to amplify synthesized AIRE to ensure compatibility with downstream steps, see Data S2 for Primers.) in turn, based on previous work^12^. Primers were also designed that ‘tiled’ across the *AIRE* coding region, splitting it into fifteen ∼150-200 base pair tiles with inclusion of overhang indexing adaptors to allow for short-read Illumina sequencing. The adaptors used for overhangs were the standard Illumina-compatible TACACGACGCTCTTCCGATCT overhang for the F primer and AGACGTGTGCTCTTCCGATCT for the R primer. We also inserted 1-3 “N” nucleotides between the adaptors and the coding sequence for each tiling primer, to increase sequencing library complexity by offsetting the nucleotide sequenced for each read. See Data S2 for a list of all mutagenic and tiling primers used. Generation of the mutagenic library via DT-POPCode was accomplished by hybridizing mutagenic oligos to each one of three WT *AIRE* mutagenic regions (Region 1: amino acids 1-181; Region 2: amino acids 182-362; Region 3: amino acids 363-545) while simultaneously hybridizing two oligos (the “dual tags” that give DT-POPCode its name) that allow primer extension from the 5’ end and (after ligation) addition of a 3’ end that enables us to later specifically amplify the mutagenized product using PCR. We performed quality control of the linear amplicon library using Illumina sequencing of tiles, and counted variants with the TileSeq_MutCount pipeline (https://github.com/rothlab/tileseq_mutcount). A suitable coverage of >85% of substitutions present at a frequency above 10 counts per million reads and an acceptable average number of amino acid substitutions per clone (λ) between 0.4 and 1, was achieved for each mutagenic region. Next, for each *AIRE* mutagenic region, we first carried out large-scale BP cloning of the linear amplicon library into pDONR223 and transformed the product using NEB^®^ 5-alpha Competent (High Efficiency) *E.coli* and selected using LB plates with 1X Spectinomycin. DNA was extracted using a Midiprep, and afterwards, the product was large-scale LR Gateway cloned into pDEST-HC-REC-v2. Large-scale LR product transformations were performed using NEB® 10-beta electrocompetent E. coli (High Efficiency) and clones selected with 1X Ampicillin. To prevent variant coverage bottlenecks during cloning steps, we estimated the number of E.coli colonies expected to achieve 50 independent clones per codon substitutions as: 63 (number of possible codon substitutions based on NNN degenerate code) ✕ 182 (number of amino acids per mutagenic region) ✕ 50 (independent clones), divided by λ. The final mutagenic libraries for each region of AIRE in pDEST-HC-REC-v2 were prepared using an endotoxin-free DNA extraction kit, and quality checked similarly to the linear amplicon library above.

### Large-scale reporter assay

The final LR library plasmid pool for each *AIRE* region was transfected into the reporter cell line using a ratio of library:Bxb1 recombinase of 10:1 by mass using FuGENE^®^ 6 (using Opti-MEM® reduced-serum medium for the transfection mix), 30 minutes after plating 15 million insulin promoter-GFP reporter cells in 10% FBS and DMEM without the presence of PS. Two biological transfection replicates were performed per *AIRE* region. The transfection was stopped after 24 hours by replacing media with 10% FBS, DMEM and 1X PS. Cells were passaged every 2-3 days at 80% confluence. At day 7 post-transfection, cells were enriched for those with integrated *AIRE* by gating on single, viable cells expressing the integration marker mCherry, and lacking the expression of BFP from the landing pad, using the SONY MA900 cell sorter. 2 million cells with integration events were sorted per regional biological replicate. Integrated, sorted cells were expanded for 7 days and the top 10% of GFP-expressing cells were sorted to enrich for functional AIRE variants. A control population of ‘non-select’ cells that were not enriched based on reporter activity were obtained separately by selecting for single viable cells that contained the integration mCherry marker (and lack of BFP) on the same day as the selection sort. Cells were expanded, pellets of 15M cells were collected for select and non-select conditions, and DNA extracted. To sequence either select or non-select cell populations, the *Bxb1* site was first amplified from the cells with primers Ef1a-Bxb1Amp_F1 and BxbAmp_R1 (see Data S2 for Primers) using 1000 ng of DNA used per one of 15 PCR reactions, and a total number of 25 PCR cycles. Final PRCs were pooled, gel extracted, and tiling of large-scale screens was performed using 10ng of Bxb1 PCR product for each tiling reaction to prevent bottlenecking of variants. Each condition had three tiling PCR technical replicates. The WT reference sequence used as a control for sequencing error was the tiled *AIRE* WT plasmid backbone sequence. Illumina sequencing for the large-scale assay was performed to a minimal depth of 1M reads/tile.

### Functional score calculation

Sequencing reads were processed as previously described^7,13,14^, by counting variant instances and translation to amino acids, using the TileSeq_MutCount pipeline (https://github.com/rothlab/tileseq_mutcount). Counts for each amino acid change were next processed using the TileSeq_Mave pipeline (https://github.com/rothlab/tileseqMave), which calculates the enrichment or depletion of variants between the non-select and select conditions. Variants were included if they reached a minimum threshold of 10 counts per million sequencing reads in the non-select condition. WT control variant instances were subtracted from the enrichment rations. Scores for each codon-level amino acid change were averaged, and standard error calculated based on the number of measurements. This analysis was performed separately for each biological replicate, and each replicate was separately re-scaled based on the calibration of nonsense and synonymous variants set to medians of 0 and 1, respectively. Biological replicate scores were averaged, and standard error propagated using the delta method. For downstream estimation of log likelihood ratios of pathogenicity (LLR_p_) and evidence levels (described below), only variants that were present in both biological replicates were included. Otherwise, for calculation of median scores by position and all other analysis, variants were included if only detected in one of the biological replicates and existing score and standard error measurement used. Functional scores are available in Data S2, noting the presence of the variant in one or both replicates, and whether the substitution measured is single nucleotide variant (SNV)-accessible, based on the original nucleotide sequence of AIRE (non-codon optimized).

### Structural analyses

The protein structure for AIRE was predicted via AlphaFold 3^15^ using the AIRE amino acid sequence and the addition of 4 Zn^2+^ ligands. The modeled structure was colored corresponding to median AIRE positional functional scores using the tileseqMave (https://github.com/rothlab/tileseqMave) colorize script^7^. The PDB structure 2KE1 was used for the PHD1 domain in complex with H3K4me0^16^. Accessible surface area (ASA) measurements for AIRE residues were obtained from FreeSASA^17^, based on the AIRE AlphaFold 3 model. Buried residues were defined by having accessible surface area (ASA)<50 Å², while surface residues were defined by ASA>110 Å² (Positions summarized in Data S2). We predicted interacting residues with PRODIGY^18,19^ using the AlphaFold 3 predictions for monomeric, dimeric and trimeric AIRE. Interacting residues were defined as those within 5.0 Å of each other.

### Variant reference sets and benchmarking

ClinVar reference set variants and gnomAD v4.1.0 allele frequency data were accessed on Aug 25th, 2025. ClinVar reference sets were filtered on one star clinical classifications. To evaluate the performance of the map on ClinVar reference set variants compared to computational variant predictors, we assessed scores using balanced precision-recall analysis as described previously^20^. Recall was calculated by evaluating the proportion of pathogenic variants correctly identified as damaging. Precision assessed the proportion of variants known to be pathogenic below a given threshold. “Balanced” refers to the use of a Bayesian method for adjusting precision to reflect what would be expected given a balanced (50%/50%) reference set of negative (benign or likely benign) and positive (pathogenic or likely pathogenic) variants^21^. The intersection of variants that were measured in the map and all computational predictors assessed was used to evaluate performance (9 negative and 28 positive variants). Although we did not explicitly exclude variants within the PHD1 domain, we note that this intersection (and therefore our performance evaluation) did not include any reference variants at positions within the PHD1 domain.

### Log likelihood ratios of pathogenicity calculations

To determine the log likelihood ratio of pathogenicity (LLR_p_) for each variant, we first defined a reference set of variants outside of the PHD1 domain that had been scored and which received at least one ClinVar review star. This yielded 16 negative and 30 positive variants (the additional excluded PHD1 domain variants included 1 negative and 1 positive variants). The score distributions for both negative and positive variant sets were estimated using probability kernel density functions based on the Epanechnikov kernel^22^, with bandwidths determined using the Scott’s rule of thumb method^23^. The LLR_p_ values corresponding to each variant’s score were each calculated as ratio of the probability densities at that score from the positive and negative distributions, respectively. Next, we used a previous calibration approach^13,24,25^ to convert LLR_p_ scores to evidence strength levels used in current variant classification guidelines^5^. Variants with scores falling outside the observed range of scores for reference variants (i.e, those with scores < -0.73 or > 1.70) were assigned undetermined evidence strengths. See Data S2 for the estimated evidence strength levels for each measured substitution.

### Cohort analyses

We assembled an APS-1 cohort from literature reports of well-characterized APS-1 cases involving *AIRE* missense variants on at least one allele (Data S2). Altogether, 98 individuals were identified, representing 20 different nationalities across four continents^26–54^. Twenty-one people were homozygous for AIRE missense variants while 45 were compound heterozygous for a missense variant and a truncating variant (i.e nonsense, frameshift, splice site). For 32 people, monoallelic missense variants had been reported suggesting autosomal dominant inheritance. A total of 50 unique missense variants were collected. Among these, 16 had not been reported to ClinVar at the time of writing, and these were formally classified in light of the *AIRE* map. All variants were classified according to the American College of Medical Genetics and Genomics/Association for Molecular Pathology (ACMG/AMP) sequence variant guidance^5^, supplemented with recommendations from ClinGen (https://www.clinicalgenome.org/working-groups/sequence-variant-interpretation) and the UK Association for Clinical Genomic Science (ACGS) (https://www.acgs.uk.com/media/12533/uk-practice-guidelines-for-variant-classification-v12-2024.pdf). The evidence from functional data was assigned strength according to recommendations by Brnich et al^55^, but was capped at a maximum of strong in both pathogenic and benign directions.

UK Biobank analysis was in accordance with the associated data use agreements, with consent to participate having been obtained via the UK BioBank. Our use of de-identified data was approved and overseen by The UK BioBank Ethics Advisory Committee (Project ID: 51135). ICD10 codes used for UK BioBank analyses are as follows: insulin-dependent diabetes mellitus E10, adrenal insufficiency E27.1, hypoparathyroidism E20, hypoparathyroidism and other E21, candidal stomatosis B37.0, chronic atrophic gastritis K29.4, vitamin B12 deficiency anemia D51, coeliac disease K90.0, keratitis H16, gastro-enteritis and colitis K52.9, interstitial pulmonary diseases J84, seropositive rheumatoid arthritis M05, systematic sclerosis M34, thyroiditis E06, vitiligo L80, and chronic hepatitis K73. We defined “damaging” variants for each of the two computational predictors (REVEL^56^ and AlphaMissense^57^) using thresholds provided by the authors (REVEL: scores > 0.5, AlphaMissense: “pathogenic” annotations).

## Results

Here we undertook missense variant effect mapping for AIRE (outlined in Figure 1A). At a high level, this involved development and validation of a reporter assay, optimization and application of a mutagenesis strategy, identification of variants that were enriched and depleted within reporter-expressing cells. Biochemical analyses agreed with expectations and identified novel positions important for AIRE function. We also describe AIRE variants that significantly increase transcriptional reporter activity and follow up on these and novel loss-of-function variants to validate these new findings. Benchmarking of the map against a set of clinical reference variants allowed us to provide functional evidence for 242 current VUS. Additionally, we identify a genotype-phenotype correlation for scores and APS-1 severity for an international patient cohort, and associate decreased-function variants with hypoparathyroidism in a dominant model of inheritance in the UK BioBank.

**Figure 1:**
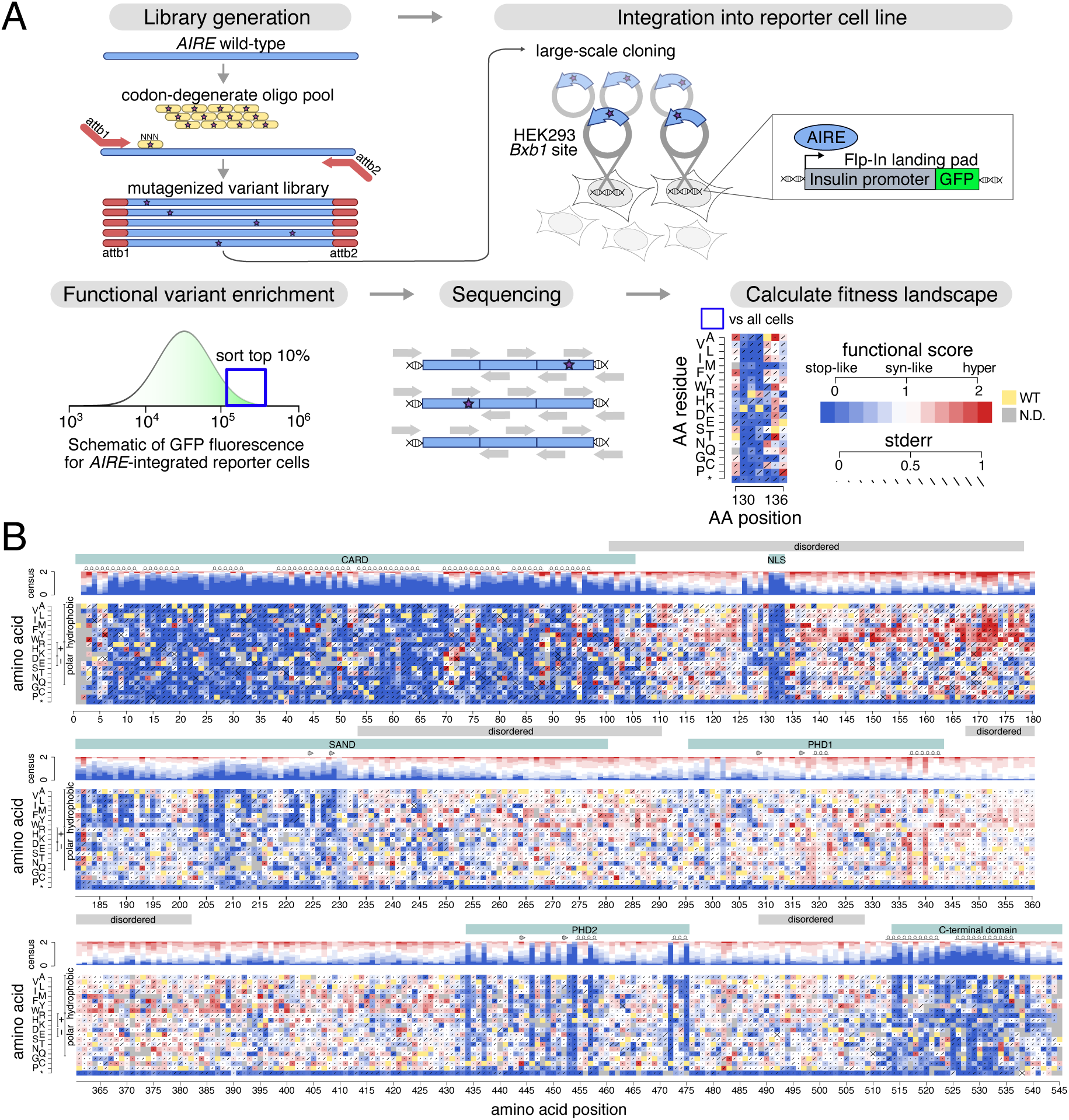
Generation of variant effect map for *AIRE*. A) Overview of variant effect mapping process for *AIRE*. B) *AIRE* variant effect map, with scores generated via an integrated insulin promoter-driven GFP reporter in HEK293 cells, and with *AIRE* variant constructs stably integrated and expressed at the *Bxb1* landing pad. WT residues at each position are in yellow, functionally deleterious variants are in blue, synonymous-like variants are in white and variants exhibiting increased reporter expression (hyper-activation) are in red. Standard errors (stderr) for each variant are represented by diagonal bars centered on each missense variant’s square. A consensus track above each position summarizes scores at that position. Domains are indicated in teal, regions of disorder in grey, and secondary structure is indicated via arrows for beta strands and helices for alpha helices.

### 2.1 The insulin promoter-GFP reporter recapitulates known AIRE variant effects

A key element of every variant effect mapping project is the development of a reliable and scalable assay that measures a molecular function of the target gene or element. Given the clinical relevance of AIRE, we also wished to ensure that we measured a disease-relevant AIRE function, namely the ability of AIRE to induce expression of genes that would not otherwise be expressed. It was therefore important to select a suitable AIRE-induced gene to assay missense variant effects. Transcriptional reporter assays in the literature have been described for AIRE, but none of these assays were immediately amenable to a high throughput approach to evaluate AIRE function. Many of these assays used the insulin promoter sequence upstream of luciferase as a reporter^58–63^, or evaluated endogenous insulin upregulation by AIRE^61,64^. As insulin is one of the thousands of genes upregulated by AIRE, because dysregulation of insulin is known to lead to either insulin tolerance or autoimmunity^60^, and due to insulin autoantibodies being implicated in the development of T1D, a symptom of APS-1^65–67^, the insulin promoter was chosen as the target to evaluate AIRE function.

Although AIRE’s role in removing or redirecting effector T cells that recognize self-antigens is thought to primarily be in mTECs, HEK293 cells have become accepted as a reliable model for the study of AIRE function, and indeed many of AIRE’s interactions in mTECs are recapitulated in HEK293 cells^64,68–73^. Therefore, we established the reporter system in a HEK293 cell line. More specifically, we used a derivative of the HEK293 cell line with two ‘landing pads’ that are useful for this study: The first contains a Bxb1 recombination site that allows for efficient large-scale genomic integration of a target construct^7,8,74,75^, enabling the generation of heterogeneous cell pools in which each cell expresses a single specific variant of the target gene. The second landing pad is a Flp-In site enabling integration of our reporter construct consisting of an insulin core promoter (previously shown to be sufficient for insulin expression^76^) fused to a GFP fluorophore.

To evaluate the reporter system, site-directed mutagenesis was used to generate two test sets of AIRE variants, with the first containing variants with pathogenic or likely pathogenic (henceforth ‘pathogenic’) annotations from ClinVar and the second containing variants with benign or likely benign (henceforth ‘benign’) annotations. These variants were stably and clonally integrated into the Bxb1 site of the insulin-promoter GFP reporter cell line. Variant function was evaluated using flow cytometry: functional AIRE was expected to upregulate the insulin promoter-GFP reporter more than functionally abnormal variants of AIRE. In cells bearing pathogenic variants, there was a clear downward shift in GFP levels (Figure S2). Although the GFP distributions of cell populations carrying pathogenic and benign variants show some overlap, Figure S2 shows that use of a specific GFP gating threshold yields clear differences between pathogenic and benign variants in terms of the fraction of cells exhibiting expression above that threshold. Therefore, the reporter system demonstrated concordance with expected variant effects and was amenable to high-throughput fluorescence-activated cell sorting for evaluation of AIRE variants at large scale.

### 2.2 Large-scale library generation for AIRE

To generate a large-scale clone library, ideally covering all possible single-amino acid substitutions in AIRE, we developed Dual-Tag Precision Oligo-Pool based Code Alteration (DT-POPCode) which eliminates a requirement of the previous POPCode approach^7^ for a uracilated template (see Methods). DT-POPCode was used to generate three amplicon libraries of the full-length AIRE cDNA, each focused on a different region of the protein. (Regional mutagenesis enables each variant to have a frequency within its library that is well above the background rate at which a variant appears simply due to base-calling error). For each library, all possible missense variants of AIRE within the corresponding region were generated using NNN degenerate oligos centered on each codon. Variant library coverage was confirmed with Illumina sequencing after *en masse* Gateway cloning into the integration vector, generating pools that contained 1.2, 0.96, and 0.62 mutations per clone for Regions 1-3 respectively (Figure S3A). Experience has shown that, for variants appearing less than 10 times per million, it is challenging to evaluate relative variant representation before and after a selection procedure. Even considering the variant above this threshold, our libraries collectively contained nearly all amino acid changes (98, 97 and 98% for regions 1-3; Figure S3B).

### 2.3 Systematical functional evaluation of AIRE missense variants

We next conducted large-scale transfections to introduce the corresponding integration vector library (with mutagenesis focused in that region) into cells bearing the insulin promoter-GFP reporter. These were carried out separately focusing mutagenesis in each of the three regions, with two biological transfection replicates for each of the regions. To select for cells with an integrated AIRE construct, cells were first sorted to obtain viable single cells exhibiting mCherry but not BFP expression (Figure S4). To enrich for cells bearing functioning AIRE variants, these cells were first expanded and then sorted again to retain the top 10% of GFP-expressing cells (‘select’ cells). We also obtained a control population of ‘non-select’ cells sorted to ensure at least some detectable GFP expression (indicating presence of a functioning reporter construct). After expansion, genomic DNA was extracted from both select and non-select cell pools, and PCR was used to amplify the integrated AIRE cDNA locus from each pool. To evaluate the frequency of each variant within each cell pool, we used a set of 200bp tiles that collectively covered the entire AIRE coding region in each library. These tiles were amplified using primers that introduced flanking sequences which enabled duplex Illumina sequencing. To derive a functional score for each AIRE variant, we used the TileSeq pipeline^7^. Briefly, this involved first calculating the log-ratio of each variant’s frequency in select relative to non-select cells. Log ratio scores were rescaled to functional scores (see Materials and Methods for details) such that the median functional score of nonsense variants was 0, and the median functional score of synonymous variants was 1. Further steps (see Materials and Methods) included estimating experimental error based on trends in replicate agreement as a function of variant abundance in the non-select library, filtering to keep scores only for those variants that were well-represented within the non-select library and–for each variant–taking the mean of biological replicate scores and propagating estimated experimental error to each final score (see Materials and Methods). Thus, we generated an AIRE missense variant effect map (Figure 1B) with scores for 9790 (90%) of the 10900 possible amino acid substitutions in AIRE, including 3127 (98%) of the 3189 substitutions that are achievable via SNVs, which are more likely than other amino acid changes to be observed clinically.

### 2.4 *AIRE* variant effect map quality evaluation

One initial measure of multiplexed assay quality is correlation between biological replicates for missense variants, which was moderate but significant (Pearson’s R=0.49, p<2×10^-16^; Figure S5A). Another measure of map quality is the extent to which scores for synonymous variants can be distinguished from those of nonsense variants: the score distributions of nonsense and synonymous variants were well separated (p<2×10^-16^ by Wilcoxon test; Figure S5B). Proteins can often tolerate nonsense variants at the C-terminus and indeed the last 8 amino acids appeared not to be essential for AIRE to activate the insulin promoter in our assay. Not surprisingly, separation between nonsense and synonymous variants remained significant after removal of these C-terminal nonsense variants (p<2×10^-16^ by Wilcoxon test; Figure S5C). Another test was whether scores followed the expectation that alleles that are more common in a human population would be less damaging. Indeed, the median scores of singletons (defined here by minor allele frequency (MAF) of < 1×10^-6^ in gnomAD v4.1.0) were more intolerant to variation than others (median score singletons =0.88, median scores common=0.95, p=4×10^-3^ by Wilcoxon test).

The quality of a variant effect map can additionally be assessed by how well it corresponds to general biochemical trends. Usually, buried amino acid residues are less tolerant of variation than surface residues, especially when buried hydrophobic positions are disrupted^77,78^. To compare the scores between surface and buried residues of AIRE, solvent accessible surface areas were predicted (using FreeSASA applied to the AlphaFold 3 AIRE structure^15,17,79^), and annotations of residues as either buried or surface (Figure S6; ASA<50Å², ASA>110Å², respectively) were compared to map scores. Although the predicted AIRE AlphaFold structure is of low confidence, we observed significantly more deleterious scores for predicted-buried residues than predicted-surface residues (Figure 2A, median map score at buried positions=0.43, median map score at the surface=0.89, p=3×10^-14^ (Wilcoxon), delta medians=0.46).

**Figure 2:**
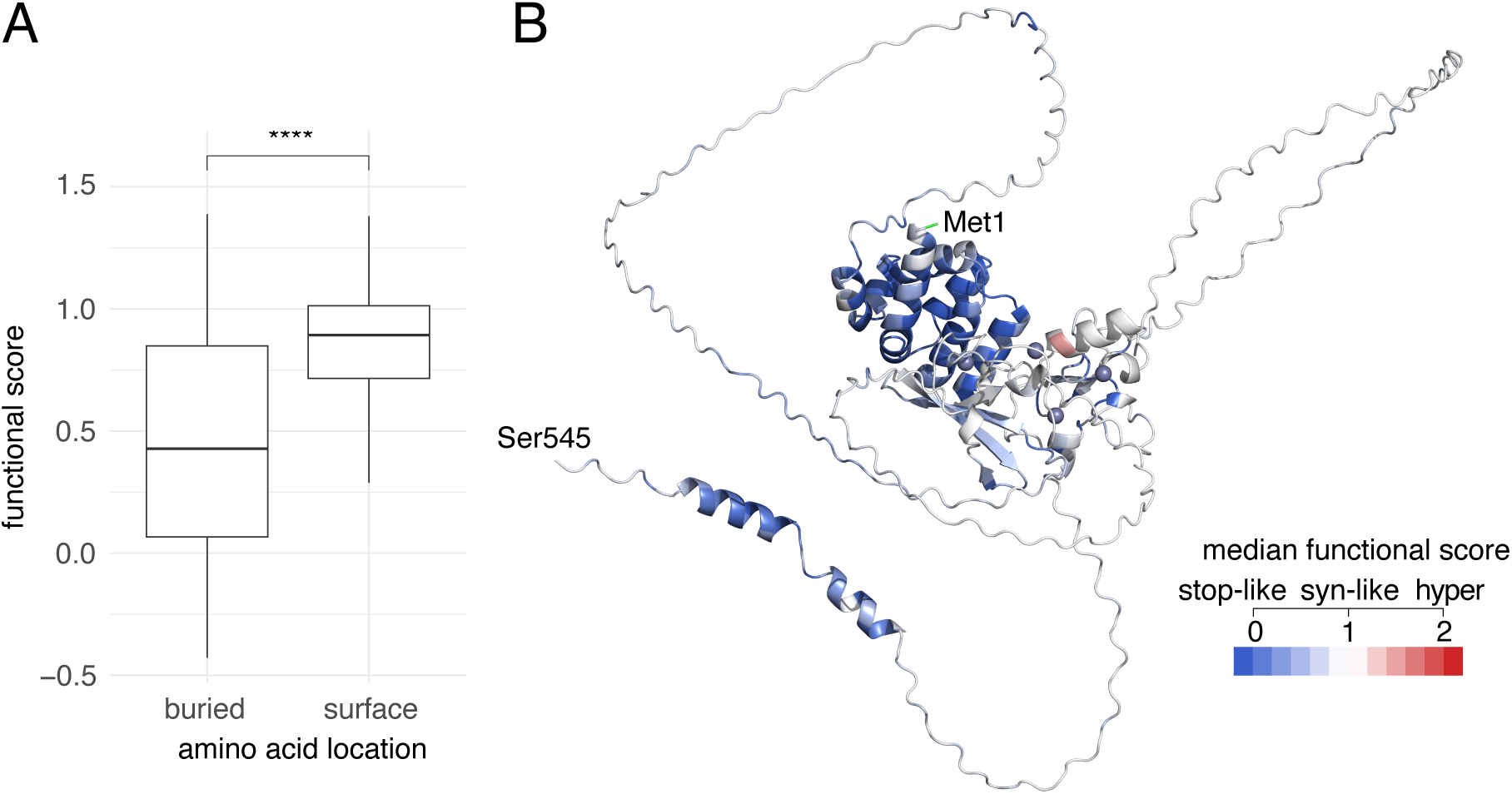
*AIRE* variant effect map scores in structural context. A) Distributions of *AIRE* functional scores corresponding to buried and surface positions. Buried and surface predictions were determined by FreeSASA on the AlphaFold 3^15^ structure prediction of AIRE (accessible surface area (ASA)<50Å² for buried, ASA>110Å² for surface). Boxes represent interquartile range, and bold bars represent medians. Annotation with “****” indicates p<0.0001 by Wilcoxon test. B) Predicted AIRE structure (AlphaFold 3) ‘painted’ according to median map scores at each amino acid position. Blue indicates positions intolerant to variation and red indicates those with apparently-increased insulin promoter-GFP reporter activity. Start and stop codons are indicated with Met1 and Ser545, respectively.

We next evaluated trends relating our map scores to the reference (“from”) and destination (“to”) amino acids (Figure S7). The lowest variant map scores were found for variants that replaced large hydrophobic amino acids (score medians for replacement of Phe, Trp, Tyr and Ile were 0.28, 0.53, 0.33, 0.28 as compared to replacement of other amino acids where medians ranged from 0.55 to 0.99). Not surprisingly, large hydrophobic residues were enriched in buried residues (representing 65% of buried positions, but only 24% of surface positions) and also in known AIRE structured domains. Other than one Tyr residue found at amino acid position 394, none appeared within predicted disordered regions.

We evaluated trends related to changing within or between different categories of amino acid such as hydrophobic, polar, and positively or negatively charged. Here we expected within-category substitutions to have a higher (less damaging) score than those between categories. Indeed, median scores for within-category substitutions ranged from 0.73-0.95, with the most damaging changes being from one positively-charged residue to another and the least damaging being from one negatively-charged residue to another (Figure S8). Median scores for between-category substitutions ranged from 0.59-0.9, with the most damaging (0.59-0.6) being substitutions of one charged residue for another with opposite-sign charge. We also found that introduction of hydrophobic residues tended to increase reporter activation (p=2×10^-13^; Figure S9A) — discussed more below (Section 2.6) — and that introduction of negatively-charged residues tended to decrease reporter expression (Figure S9B, p=7×10^-8^), while introduction of positively charged residues did not (Figure S9C, p>0.05).

We expected our map to capture the trend that introducing proline disrupts secondary structure^80^. Indeed, median scores of initially hydrophobic, polar, positively or negatively charged positions substituted to proline ranged from 0.67-0.73 (Figure S8). In general, substitutions to proline had the lowest (most damaging) median scores compared with substitutions to all other amino acids (Figure S9D, p=3×10^-4^). These trends are in keeping with conclusions from larger-scale analyses of other variant effect maps that introducing proline is generally more damaging than introducing other amino acid changes^81,82^. Interestingly, replacement of prolines by other amino acids in other categories was often tolerated, with median scores ranging from 0.77-1.07 (Figure S8).

### 2.5 Domain-based biochemical analyses reveal both known and novel AIRE features

Domains and features known to be important for AIRE function were largely (with a major exception described below) recapitulated within our map. ‘Painting’ the AIRE AlphaFold 3 predicted structure based on consensus map scores demonstrates the expected biochemical features while also suggesting novel functional features (Figure 2B).

For example, substitutions within the CARD domain (positions 1-104), which is essential for AIRE multimerization and its ability to assemble transcriptional condensates, were generally deleterious in the map as expected. Within the CARD domain, there are hydrophobic residues within the alpha-helical bundle that point into the core of the card domain according to the predicted structure, and these were found intolerant to substitution in our map (Figures S10A and S10B; median buried = 0.03, median surface = 0.22, p>0.05). Residues within the outer alpha helices of the CARD domain’s alpha-helical bundle appeared more tolerant to variation than those of the innermost helical residues. Thus the importance of the CARD domain was well-captured by our large-scale insulin promoter-GFP reporter assay.

In the predicted monomer structure (Figure 2B) CARD domain appears centrally located within the AlphaFold model, but we cannot be confident that it is buried given that the rest of the structure is of low confidence. We also generated AIRE dimer, trimer and tetramer structural models using AlphaFold 3^15^ (Figure S11), and identified residues that were buried by contacts between dimeric, trimeric or tetrameric subunits using PRODIGY^19^ (Positions summarized in Data S2). These multimerization models are supported by the observation that the residues involved in interaction showed less tolerance to substitution than non-interacting residues (Figure S11). Furthermore, the residues involved in dimerization demonstrated the greatest difference between interacting scores compared to those involved in trimerization or tetramerization (Δmedian difference score was 0.26, 0.24, 0.20 interacting and non-interacting positions for dimer, trimer and tetramer respectively), implying residues involved in AIRE dimerization are more influential on AIRE function than those involved in trimer or tetramer positions.

AIRE is required to localize to the nucleus to be able to interact with transcriptional machinery. Based on sequence patterns shared with other nuclear localization (NLS) sequences, the AIRE NLS was previously described as bi-partite, including positions 110-114 and 131-133^83^. However, a later study concluded based on deletion analysis that the NLS was monopartite, consisting only of positions 131-133^84^. In agreement with the latter study, our map (Figure 1B) showed position 110-114 to be tolerant to substitution, but intolerant to variation at positions 131-133. Thus, our results support both the conclusion that the AIRE NLS is required and that it is monopartite.

The AIRE SAND domain (positions 181-280), which has been implicated in DNA and protein-protein interaction, is divergent from other SAND domains such as those in proteins NUDR, Sp100b, GMEB1 which all contain the DNA-binding surface residues KDWK. In AIRE^85^, these residues evolved to NKAR (positions 244-247) which exhibit mixed tolerance to variation with neutral scores (median scores of NKAR positions ranged from 0.61 to 0.85). The predominant current model is that AIRE upregulates genes that contain poised transcriptional machinery at double-stranded breaks^86^, but it is not clear whether or not this requires DNA binding by AIRE (which has been proposed by some but has been controversial)^64,72,87^. This map provides modest support for the proposition that the (potentially) DNA-binding residues NKAR are not essential for upregulation of the reporter.

Within the SAND domain, the map shows intolerance to substitution with hydrophobic residues within positions 181-223, suggesting that this N-terminal segment of SAND is located on a solvent-accessible surface of the protein (Figure 1B). Together with previous evidence that SAND mediates protein interaction^88^, this result offers the possibility that this segment provides interfacial residues in SAND’s protein-protein interactions. Overlapping the C-terminal end of this segment, the beta strands of the SAND domain (positions 221-231) were the only residues of the SAND domain that appeared structured in the predicted AlphaFold structure and these appeared highly intolerant to variation in our map (Figure S12A). Interestingly, a segment just C-terminal to SAND’s beta strands (positions 234-290) that is annotated as intrinsically disordered (despite overlapping with the annotated SAND domain) did also have some positions (Phe236, Asn244, and Arg257) that were intolerant to substitution (median positional scores of 0.63, 0.60 and 0.65, respectively). The C-terminal half of the SAND domain is disordered and generally tolerant to substitution, although largely intolerant to the introduction of charged residues (Figure 1B). Overall, our map supports the importance of SAND domain in AIRE function, and that its importance may in part be attributable to engagement in protein-protein interactions.

The PHD1 domain (residues 296-343) domain, which interacts directly with H3K4me0^64^, harbours most of the known pathogenic *AIRE* variants that have an autosomal dominant mode of inheritance^52^. Previous small-scale studies have recapitulated these pathogenic effects using reporter assays^61,63,64,89–93^. However, it is unclear what we should have expected from our map in this domain, given that we did not carry out our measurements in the presence and absence of a wild-type allele, as may have been needed to detect dominant negative effects. Indeed, the median score at PHD1 positions was 0.98, indicating a high tolerance for substitution, and our map clearly did not capture pathogenic variant effects known for this domain. More specifically, scores for residues known to be directly involved in the interaction of PHD1 to H3K4me0 appeared to tolerate variation in our map (Figure S12B). That our map failed to capture the functional importance of the PHD1 domain can also be seen by painting the average score by position onto the existing NMR structure for the PHD1 domain interaction with H3K4me0, demonstrating that the H3K4me0-proximal residues appeared to tolerate modification in our map (Figure S12C). This limitation of our map at the PHD1 domain might be explained if the Flp-In site where we integrated our insulin promoter-GFP reporter construct is in open chromatin and therefore does not require a functional AIRE PHD1 domain.

Exceptions to the general observation that our map did not capture PHD1 function include residues Cys299 (median positional score = 0.56) and Cys302 (median positional score = 0.55), confirm Zn^2^-binding importance as they are directly proximal to the first Zn^2^ ion of PHD1 (Figure S12C). Thus, while the PHD1 domain does not show the expected intolerance to variation of the H3K4me0-proximal surface, Zn^2+^-bound residues still show variant effects. Another position expected to be intolerant to variation was Cys311 in the PHD1 domain, as the patient variant p.Cys311Tyr has been hypothesized to impact disease through decreased zinc-chelation^52,94^, and our results suggest that this position is likely not essential for zinc interaction. Together, these results suggest Cys311 is more important for H3K4me0 binding than zinc-binding. Interestingly, the PHD1 Cys337 and Cys340 residues showed increased reporter activity when modified (median positional scores = 1.30 and 1.39, respectively; Figure S12C), although the reasons for this are unclear.

The PHD2 domain is a zinc-finger domain known to be important for protein-protein interactions that coordinates two Zn^2+^ ions^83^. The map reveals that within the PHD2 zinc-binding domain, all zinc-adjacent cysteines are deleterious, (Cys434, Cys437, Cys446, Cys449, Cys457, Cys472 and Cys475) in addition to zinc-proximal His454 (Figure S12D). The map captures the importance of Zn^2^ binding of the PHD2 domain to transactivation of the reporter.

AIRE’s C-terminal domain (positions 514-545) has previously been demonstrated to be important for its transactivation activity, probably through protein-protein interactions with transcriptional coactivators CBP/p300 localized to enhancers^92,95^. Our map showed general agreement with this finding in that variation in this region was not tolerated (Figure 1B). However, our map showed that nonsense variation as described above was tolerated within the last 8 amino acids. An interesting exception to this trend was that the map found substitutions introducing a positively charged residue within the final 8 positions to be damaging (Figure 1B).

### 2.6 Unexpected findings from the AIRE map

Two residue positions that displayed intolerance to variation but did not correspond to known functional roles were Pro126 and Pro129 (median positional scores = 0.31 and 0.46, respectively). These fell within a disordered region from positions 101 to 178 with very low confidence of structure from according to AlphaFold^79,96^ These prolines fit with a larger pattern of many prolines in positions 115-129 (Pro115, Pro116, Pro119, Pro124, Pro125, Pro126 and Pro129). More generally, prolines in intrinsically disordered regions have been described as both disorder promoters and involved in structural compaction^97^, but only Pro126 and Pro129 were intolerant to variation, with Pro126 being especially intolerant. We found it interesting that these important prolines were close to the NLS (previously described to be at positions 131 to 133^84^). Thus, our map allows us to suggest that the function of Pro126 and Pro129 is to maintain proper AIRE conformations that are necessary for NLS function.

Many (3484) missense variants exhibited reporter activation that was numerically higher than that of wild type (i.e., greater than 1). Interestingly, visual examination of the map showed that, within disordered regions in general (excluding the NLS) and especially at SAND domain positions 145-175, greater-than-wild-type reporter activity was frequent for substitutions that newly introduced a hydrophobic residue (Figure 1B). Indeed, substitution to a hydrophobic residue within a disordered region significantly increased reporter activation (median functional score difference between substitutions to hydrophobic and non-hydrophobic amino acids was 0.14, p=1×10^-22^; Figure S13A). However, within non-disordered regions, no significant difference was found between substitution to hydrophobic substitutions and non-hydrophobic substitutions (median difference = 0.03, p>0.05; Figure S13B). This phenomenon could be explained by the fact that hydrophobic residues on surface-exposed positions typically provide much of the energy for protein-protein binding^98^, with newly introduced hydrophobic residues in disordered regions potentially providing increased binding energy.

A caveat here is that even if there are many true gain of function variants that enable us to detect convincing trends like the one just described, it is harder to be sure that any specific variant is truly gain of function. Some variants will get scores >1 simply due to experimental error (e.g., even variants with truly wild-type function would be expected to yield scores numerically higher than 1 roughly half of the time). To distinguish true ‘gain-of-function’ variants, we therefore assessed one-tailed empirical P-values for each variant using synonymous variants as the reference score distribution. In total, 8 variants were found to significantly upregulate the reporter (p<0.05 after adjusting for multiple testing; Supplemental Table 1).

Amongst variants that we observed to significantly upregulate the reporter, the three highest-scoring were p.Ala138Met, p.Pro162His, and p.Pro162Tyr. To confirm that novel map findings can be replicated, we applied a small-scale assay to these, to the two low-scoring variants p.Pro126Val and p.Pro129Arg that were discussed above as being potentially related to nuclear localization, as well as additional controls. Flow cytometry demonstrated that all variants and controls behaved in accordance with their map scores (Figure S14), with p.Pro126Val, p.Pro129Arg, and negative controls upregulating the reporter significantly less than WT. The three high-scoring variants showed reporter expression significantly above WT, representing gain of transcriptional function. Of the missense variants which numerically increased reporter expression, 35% are ‘SNV-accessible’ (i.e., the new amino acid can be encoded by changing a single nucleotide in the reference codon), and so are likely to already exist in the population. However, only three (p.Thr46Met, p.Pro162His and p.Lys395Met) of the 8 variants (Supplemental Table 1) exhibiting significantly increased reporter expression are SNV-accessible. As described below, UK BioBank analysis did not support an association between variants exhibiting increased reporter expression and altered frequency of AIRE-related symptoms. However, this analysis was based on only 45 participants who all carried the SNV-accessible VUS variant p.Thr46Met (the only variant significantly above a score of 1) that was observed in the UK Biobank, and was therefore not well powered to detect subtle effects.

### 2.7 Using the AIRE map to classify clinical variant pathogenicity

To evaluate the potential of our AIRE map in classifying variant pathogenicity, we first analyzed the tradeoffs between precision (the proportion of variants below a given score that are annotated P/LP) versus recall (the proportion of variants that are annotated as P/LP and scored below a given threshold). More specifically, we reported “balanced precision”, defined as the precision that one would have expected to observe with a balanced test set with equal numbers of pathogenic and benign reference variants, which was derived from nominal precision as previously described^21^. Here, our variant reference set was composed of 17 variants annotated as B/LB and 36 variants annotated as P/LP with at least one star in ClinVar. Of these, map scores seen in both biological replicates were obtained for 16 B/LB and 30 P/LP. We next evaluated computational predictors including SIFT^99^, ESM1b^100^, VARITY^21^, REVEL^56^, and AlphaMissense^57^. Reducing our reference set to the intersection of variants that were both scored by our map and by computational predictors yielded 9 B/LB and 28 P/LP variants (which we note did not include any variants in the PHD1 domain). A balanced precision recall curve generated using this reference set (Figure 3A) showed that the map, and most predictors, performed well. REVEL and VARITY were the best predictors in terms of recall at a fixed stringent (90%) precision (R90BP: map=1, ESM1b=0.61, AlphaMissense=0.93, VARITY=1.00, SIFT=0.29, REVEL=1.00; Figure 3A). Overall, our map performs on par with the computational predictors REVEL, AlphaMissense and VARITY in separating pathogenic from benign missense variation.

**Figure 3:**
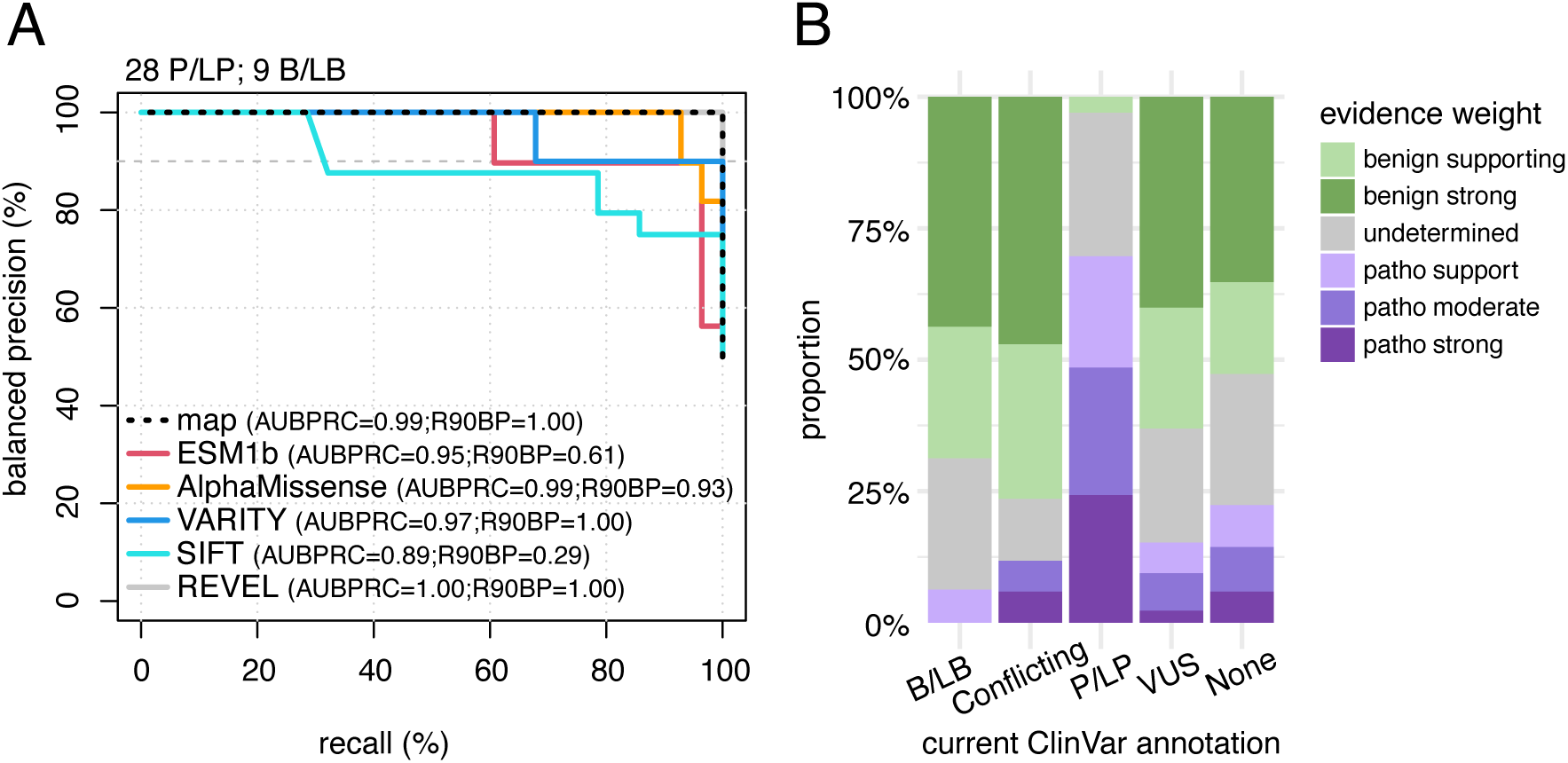
Benchmarking of *AIRE* map to assess performance. A) Performance of *AIRE* variant effect map and computational predictors of variant effect (ESM1B^100^, AlphaMissense^57^, VARITY^21^, SIFT^99^ and REVEL^56^) by balanced precision vs recall analysis based on a variant reference set scored by the map and all predictors tested (28 P/LP and 9 B/LB variants). B) Proportion of map-derived evidence labels for missense variants currently classified on ClinVar to B/LB, Conflicting, P/LP and VUS, and for single nucleotide-accessible variants with no current clinical annotation (None).

We next sought to calibrate map scores to yield a quantitative log-likelihood ratio of pathogenicity (LLR_p_) corresponding to each score. LLR_p_ scores are not only compatible with Bayesian methods for variant interpretation, but can also be converted into the categorical evidence strength labels used within the ACMG/AMP framework^13,24,25^ . Briefly, LLR_p_ scores were calculated in two steps: First, probability density distributions for B/LB and P/LP variants were separately estimated using observed map scores for each set, using kernel density estimation (Figure S15A). These distributions were estimated using reference variants in each set that had been measured in both biological replicates. Given the above-mentioned limitations of our map in detecting damaging variants in the PHD1 domain, we performed this conversion of scores to evidence strength labels using a reference set of 16 B/LB and 30 P/LP variants that excluded one P/LP and one B/LB variant in the PHD1 domain. Second, at each possible map score, LLR_p_ was calculated as the log_10_ of the ratio of P/LP probability density to B/LB probability density. As previously described^13,24,25^, LLR_p_ scores were then converted to ACMG/AMP evidence strength labels (Figure S15B). This procedure yielded non-undetermined functional evidence for 70% of variants currently annotated as VUS (Figure 3B). Although we retained evidence provided for PHD1 variants with evidence towards pathogenicity (Figure 3B), we declined to provide evidence towards benignity for variants in the PHD1 domain. As a further element of conservatism in providing clinical evidence, we added a “caution” flag to evidence where the confidence interval of score overlapped 0.5 (the midpoint between the medians of nonsense and synonymous variants; caution flags are summarized in Data S2).

We next evaluated concordance of our functional evidence with current annotations of our reference set variants. Our new evidence was concordant with classifications for 39 (89%) of 44 variants if PHD1 variants are excluded and 40 (87%) of 46 if PHD1 variants are included (Figure S16). Of the 6 variants for which the LLR_p_ pointed in the opposite direction from current annotation (Supplemental Table 2), four were not truly discordant in the sense that our map provided no evidence strength label (in one case because we declined to provide evidence towards benignity for PHD1 domain variants). Of the two discordant variants, one was p.Arg9Pro, a known pathogenic variant to which we assigned supporting evidence towards benignity, but this variant received a caution flag based on its confidence interval. The other was p.Arg471Cys, a common variant (MAF of 1-2% in Northern Europeans) which is annotated as benign but which our map assigned pathogenic supporting evidence. That p.Arg471Cys is a known contributor to polygenic risk for Addison’s disease, pernicious anemia and type 1 diabetes^101–105^, supports the validity of our map. Overall, our assay largely recapitulated expected reference set variant effects, and we generated evidence for 6103 of AIRE variants (of which 1019 had a caution flag) that do not currently have a clinical classification (Figure 3B).

We next re-evaluated the pathogenicity of all *AIRE* missense variants in ClinVar in light of new evidence from our map, using ACMG/AMP criteria as described in Materials and Methods. Variants were subjected to re-evaluation if our map offered a calibrated evidence strength label. Of the 13 conflicting variants which were VUS/B/LB conflicting, 10 were re-classified to B/LB, and 3 remained as VUS. Of the 2 conflicting variants (for which both a VUS and an LP classification had been reported on ClinVar), one was assigned P both with and without our evidence, while the new evidence led us to reclassify the other from VUS to LP. Among the 242 currently-VUS variants evaluated, 133 (55%) were re-classified: 14 (6%) to P/LP (13 to LP and 1 to P) and 49% (119) to LB. Of the 12 B/LB and 23 P/LP we re-evaluated, all remained in the same category except the current P/LP p.Arg9Pro which was downgraded to a VUS albeit with low confidence (caution flag). Thus, evidence from our largely recapitulated previous pathogenicity annotations, while enabling re-classification for a total of 133 variants which had previously been classified as VUS (109 of which were of high confidence, see Data S2).

### 2.8 Dominant negative variant impacts

Some AIRE variants exhibit dominant inheritance, and are thought to be dominant negative (DN) as a result of their ability to sequester WT AIRE^52,106–108^. Our assay does not convey any information about the ability of AIRE variants to sequester WT AIRE (HEK293 cells do not express WT AIRE^64^), but does allow us to assess the intrinsic activity of AIRE variants. Therefore, we evaluated map scores of a set of 18 known DN variants that have been confirmed in small-scale assays^109^, comparing them both to B/LB variants and to all other (presumed AR) P/LP variants reported in ClinVar (Figure S17A). While AR P/LP variants showed more damaging scores than B/LB variants as expected (Δmedian score 0.74, p_adj_: 8×10^-6^), non-PHD1 DN variants had scores not significantly different than those of AR variants (Δmedian score 0.47, p_adj_>0.05) or B/LB variants (Δmedian score 0.27, p_adj_>0.05). However, most known DN variants are located in the PHD1 domain, for which our assay showed reduced sensitivity (Figure 17B). Four known DN variants fall outside of the PHD1 domain. Two of these, p.Pro368Ala and p.Gly467Arg, did not exhibit damaging scores. Interestingly, the two remaining, well-characterized DN variants that fell outside of the PHD1 domain (p.Gly228Trp and p.Cys446Gly) were both found deleterious in our map, suggesting that they have a functional impact on transcription beyond sequestration impacts.

Usually, DN variants have less-severe autoimmune symptoms and are associated with organ-specific autoimmunity^52^. However, some DN variants, such as p.Cys302Tyr and p.Cys311Tyr (which are both within the PHD1 domain at positions involved in coordinating Zn^2+^-binding), are known to more closely resemble classic APS-1 symptoms^52^. Of these, p.Cys302Tyr showed moderate functional impact in our reporter assay (score = 0.47), while p.Cys311Tyr was not found to be as damaging in our assay (score = 0.79). Interestingly, a range of patient phenotypes (from asymptomatic to severe) has been reported for p.Cys311Tyr^52^. Overall, while the map has reduced sensitivity to detect DN variant impacts, we do detect the impacts of some DN AIRE variants on insulin promoter activity.

### 2.9 Comparing map scores to an international APS-1 patient cohort supports a genotype-phenotype relationship for APS-1

To evaluate the potential predictive value of our AIRE map in the context of human phenotypes, we examined a well-phenotyped cohort of 98 APS-1 patients published in the literature, including 21 with homozygous missense variants, 45 having a missense variant in trans with a nonsense or splice site variant, and 32 carrying a single heterozygous missense variant. Within this cohort, 45 unique symptoms were collectively described (Supplemental Table 3), with only 26% of these seen in more than 10 participants. A total of 16 AIRE variants that were novel (relative to ClinVar) were present in this cohort.

An initial barrier to quantitatively evaluating genotype-phenotype associations in the context of our maps was that our map provides functional information for single-variant AIRE alleles, rather than diploid AIRE genotypes. We therefore derived an combined-allele map (CAM) score by combining map scores for AIRE variants from both alleles (using simple addition of variant effect map scores for variants on different alleles, as was previously shown to yield significant genotype-phenotype correlation for the protein CBS ^12^). For the purpose of calculating the CAM score, splice site and nonsense variants were modeled as null (i.e., given scores of 0) and WT alleles were assigned a score of 1.

A second barrier to this analysis was the lack of widely accepted quantitative measures of APS-1 disease severity. To address this, we derived a simple proxy for severity: the number of symptoms reported for each patient. The number of symptoms correlated negatively with CAM scores, such that people with a lower (more damaging) CAM score tended to exhibit more symptoms (R=-0.4; p=4.5×10^-5^; n=97; Figure S18A). We worried that this correlation might have resulted from the fact that PHD1 variants may both yield less severe disease and also be less readily detected in our assay. However, after eliminating any participants with at least one PHD1 variant, a significant negative correlation remained (R=-0.29; p=0.01; n=71, Figure 4A). These results provide the first statistically significant evidence of quantitative genotype-phenotype relationship for APS-1.

**Figure 4:**
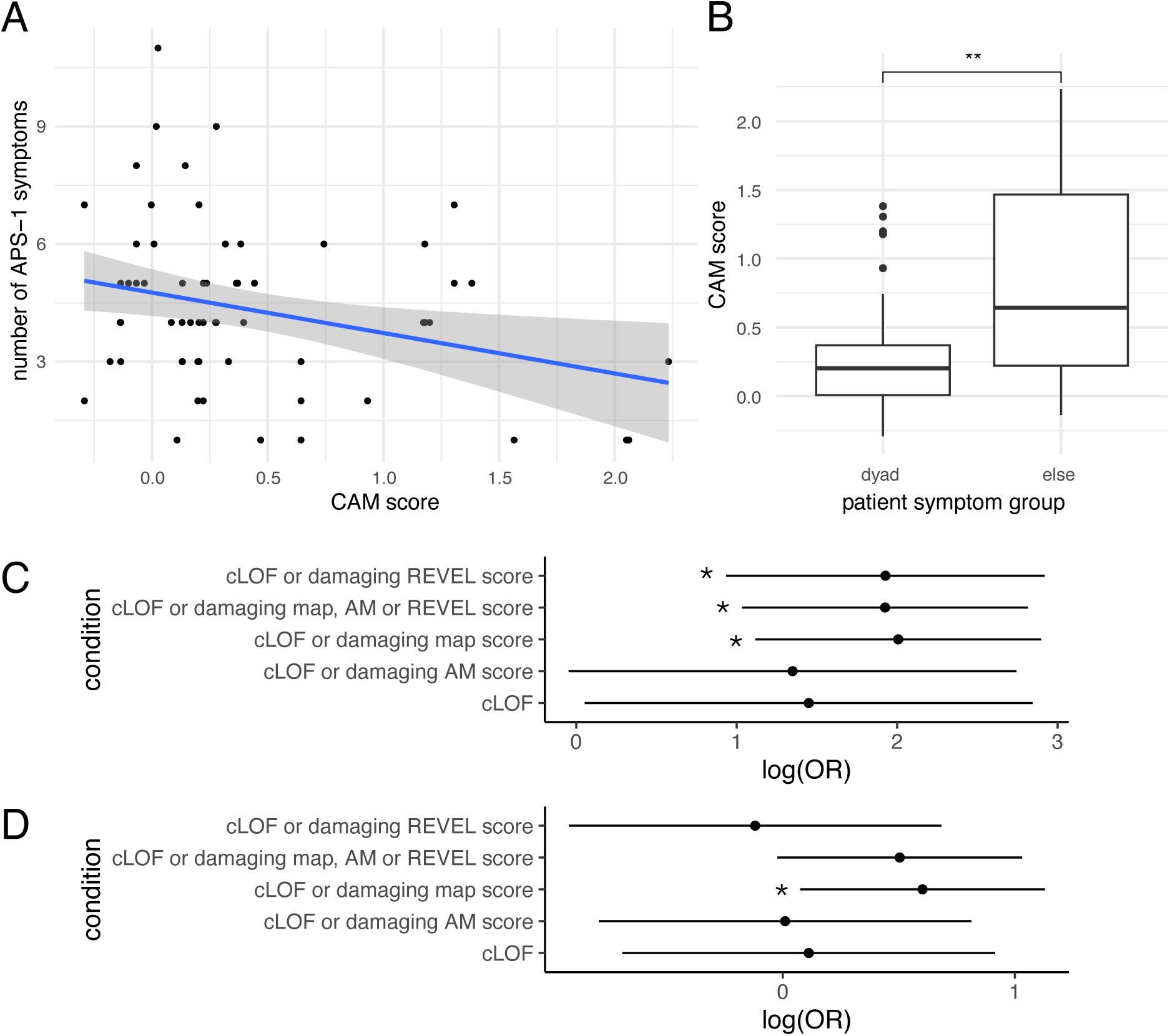
Genotype-phenotype assessment of *AIRE* map against patient cohorts. A) For an international cohort of APS-1 patients (n=71; excluding those carrying PHD1 variants in the PHD1 domain), the relationship between the combined-allele map (CAM) score (defined as the sum of functional scores for both patient alleles, where nonsense/frameshift and WT alleles are assigned scores of 0 and 1, respectively) and the number of APS-1 symptoms (R=-0.29, p=0.01). (See Figure S18 for a PHD1-inclusive version of this plot with similar conclusion.) B) Distributions of CAM scores for patients in the international patient cohort who either do (n=53) or do not (n=18) exhibit the diagnostic dyad of APS-1 symptoms (“**” indicates p=6×10^-3^, by Wilcoxon). Here we excluded patients with variants in the PHD1 domain (see Figure S18 for a PHD1-inclusive version of this figure with similar conclusion). C) Confidence intervals for the log-odds ratio of having a hypoparathyroidism diagnosis for those UKBB participants carrying at least one rare (<0.0001 gnomAD minor allele frequency) *AIRE* variant that was deemed intolerant to variation by some definition of intolerance. Here log-odds ratio = 7.4[3.1,18.1], p_adj_=7e-4 for the variant effect map and cLOF; OR = 6.9[2.5,18.5], p_adj_=3×10^-3^ for REVEL and cLOF; log-odds ratio = 6.9[2.8,16.7], p_adj_=1×10^-3^ for all predictors and map combined. D) As in C), but considering the log-odds ratio of having a Vitamin B12 deficiency anemia diagnosis. Here, log-odds ratio = 1.8[1.1,3.1], p_adj_=4×10^-2^ for the variant effect map and cLOF. Annotation with “*” =p_adj_<0.05.

As another proxy for disease severity, patients were grouped into those with and without the presence of the APS-1 diagnostic dyad (two out of hypoparathyroidism, adrenal insufficiency or chronic mucocutaneous candidiasis). Patients presenting with the diagnostic dyad had CAMs that scored significantly lower in the map than non-dyad patients (median dyad CAM=0.22, median non-dyad CAM=1.68; p=2×10^-7^; Figure S18B). Similarly, excluding patients with PHD1 variants from analysis demonstrated significantly lower CAMs for patients with the diagnostic dyad (median dyad CAM=0.20, median non-dyad CAM=0.64; p=6×10^-3^; Figure 4B). Overall, our results provide evidence for a genotype-phenotype correlation for APS-1 in this patient cohort.

### 2.10 UK BioBank participants with damaging AIRE variants associate with hypoparathyroidism

We next evaluated whether variants identified in the AIRE map were associated with APS-1 and its related symptoms in whole exome sequencing data from 469,765 UK BioBank participants^110^. APS-1 is a very rare disease, and indeed only nineteen participants had an APS-1 diagnosis in UK-BioBank at the time of our analysis, of which only three participants each had one missense variant. The lack of sequenced variants in this group could be attributed to the GC-rich content of *AIRE*, which is difficult to exome sequence.

Heterozygous *AIRE* genotypes have also been associated with autoimmune disease also in the heterozygous state^52^. To evaluate whether having at least one damaging *AIRE* allele is associated with APS-1-related symptoms we examined symptoms commonly associated with *AIRE* variants, including both endocrine symptoms (insulin-dependent diabetes mellitus, adrenal insufficiency, and hypoparathyroidism) and non-endocrine symptoms (candidal stomatosis, chronic atrophic gastritis, vitamin B12 deficiency anemia, and others (see Materials and Methods)).

We assessed the association between these APS-related symptoms in UK BioBank and patient sets defined by presence of at least one putatively damaging AIRE variant from five non-exclusive definitions of damaging: 1) classical predicted LoF (cLoF; stop or frameshift or canonical splice site variants) only, 2) either cLoF or a damaging score in our variant effect map, 3) cLoF or a damaging AlphaMissense score, 4) cLoF or a damaging REVEL score and 5) Any, i.e., cLoF or AlphaMissense or REVEL or our map (see Materials and Methods for the thresholds that define damaging variants from each score source).

Hypoparathyroidism showed consistent association with *AIRE* scores across the sets with the strongest associations coming from the variant effect map (OR = 7.4[3.1,18.1], p_adj_=7×10^-4^; Figure 4C) and REVEL (OR = 6.9[2.5,18.5], p_adj_=3×10^-3^). A combination of scoring methods (generating patient set 5, above) also showed a strong hypoparathyroidism association (OR = 6.9[2.8,16.7], p_adj_=1×10^-3^). The most conservative model (scoring only cLoF variants) captured fewer variants and did not meet nominal significance (OR = 4.3[1.1,17.2], p_adj_=0.08). We also observed increased risk for vitamin B12 deficiency anemia using a combination of cLoFs and our map (patient set 2; OR=1.8[1.1-3.0], p_adj_=0.04; Figure 4D). These results provide additional support for the role of AIRE in autoimmune disease susceptibility, and for the value of our variant effect map in identifying variants that contribute to disease susceptibility.

## Discussion

Prior to this study, experimental functional impact data had been reported for fewer than 7% of clinically-reported *AIRE* missense variants. Where they had been studied, the data often became available years after patient presentation. This lagging reactive pace of functional evaluation is unfortunate, especially given that essentially all possible single-nucleotide variants already exist in someone alive today^111^. Here we provide a systematic missense variant effect map of *AIRE*, evaluating 9790 AIRE amino acid substitutions for their ability to enhance expression at the insulin promoter. Amongst these, we assayed 3127 (98%) of the 3189 missense variants that can occur via a single-nucleotide change from the reference sequence (the amino acid substitutions most likely to be clinically observed). Importantly, this *AIRE* missense variant map provides scores proactively for yet-to-be-observed clinical variants.

Our *AIRE* variant effect map was well aligned with expectation in the sense that residue positions known to be important for structure and function were less tolerant to variation. Such positions included buried residues, NLS residues, and residues of known domains critical for AIRE function such as the CARD, SAND, PHD2 and C-terminal domain. One exception was substitutions in PHD1 domain positions which, except p.Cys299 and p.Cys302 residues involved in zinc-binding, appeared to be generally tolerated. Given that this domain is known to be important for AIRE function where promoters fall within closed chromatin, we consider this a limitation of our map. We attribute this limitation to the fact that our insulin promoter reporter construct was located within an open-chromatin safe harbor site, so that AIRE’s PHD1 function was not necessary to enhance reporter expression. Our use of an open-chromatin reporter likely provided a more robust signal that represented a compensating advantage for non-PHD1 functions, in that AIRE is known to provide expression at closed chromatin only stochastically^112^ with presumably less-robust signal output.

We used our map to identify AIRE positions, which had not been previously reported as functionally important, as having either loss- or gain-of-function impacts according to our reporter activation assay. For example, the two residues p.Pro126 and p.Pro129 were intolerant to most substitutions. Although we do not know the mechanism for the importance of these residues (and no clinical variants at these positions have been reported), they are located in a disordered region proximal to the NLS. Given that prolines have often been implicated in maintaining structural compaction of disordered regions^97^, we hypothesize that these prolines serve to maintain proper NLS conformation. Follow-up subcellular localization studies of substitutions could help understand their role. We also noted an interesting trend that variants increasing reporter activation were often hydrophobic residues introduced to disordered regions. Indeed, in small-scale follow-up testing, we were able to experimentally confirm increased reporter activation for three of these substitutions: p.Ala138Met, p.Pro162His, and p.Pro162Tyr.

The map achieved excellent performance in the sense that the score distributions for known pathogenic and benign were markedly different, especially after excluding the PHD1 domain (based on limitations discussed above). The ability of our map to distinguish pathogenic and likely-pathogenic variants from benign and likely benign variants was on par with the current standard of computational effect predictors including REVEL, AlphaMissense, and ESM1b. Current clinical guidelines^5^ treat computational and experimental evidence as independent, so that these approaches are complementary rather than being in competition. In converting functional scores to LLR_p_ scores that reflect ACMG/AMP categorical evidence strengths, we provided functional assay evidence for 71% of all SNV-accessible missense variants, including 70% of variants currently annotated as VUS, including SNV-accessible variants within the PHD1 domain that showed evidence towards pathogenicity. In addition, we provide confident, formal re-classification for 133 VUS AIRE variants according to current ACMG/AMP guidelines.

The *AIRE* map enabled us to characterize genotype-phenotype relationships within APS-1 patient cohorts. Patient-level CAM scores derived from our map correlated with two different proxy measures of APS-1 severity, and also with the presence/absence of the APS-1 diagnostic dyad. Using the UK BioBank, additional associations were found between the presence of a rare and damaging AIRE variant (i.e., using a dominant inheritance model) with both hypoparathyroidism and vitamin B12 deficiency anemia. These results complement previous reports describing mono-allelic effects for *AIRE* mutations within the PHD1 domain^52^, in that the both of the associations detected here using our map-derived CAM scores were entirely driven by non-PHD1 variants.

Given that the HEK293 cell line we used does not recapitulate all physiological conditions of the thymus *in vivo*, and given that AIRE may function differently at different target gene loci, perhaps we should have expected our assay to be uninformative about pathogenicity. If that had been the case, we might have pursued a potentially-more-faithful thymic cell line, and sought to optimize high-throughput transfection and evaluation via flow cytometry. Fortunately, as indicated in our small-scale evaluation of the HEK293-based insulin reporter assay, and as borne out in the final map, the more facile HEK293 assay appears highly predictive of pathogenicity. Success with HEK293 cells is consistent with their previous use in a range of biochemical experiments fundamental to the characterization of AIRE function^87,113^.

One limitation is that our assay was based on variant *AIRE* cDNAs, such that impacts of SNVs (either coding or non-coding) on splicing will not have been captured. In future work, by introducing *AIRE* variants at the endogenous locus and comparing the results with our cDNA-based map, one might hope to identify variant impacts on splicing, mRNA stability, enhancer and other non-coding functions.

Another limitation is that, while our assay can detect coding variant impacts on AIRE’s transcriptional function, it cannot inform us as to the mechanism of functional impact, e.g., loss of stability or weakened protein interaction, etc. Variant effect maps examining additional ‘subfunctions’ could address this, e.g., by systematically testing AIRE variant impacts on protein abundance^74^ or nuclear localization^114^.

It would be ideal to expand this work to assess other AIRE-induced genes to evaluate whether there are promoter target-specific differences in *AIRE* variant impacts. Interestingly, AIRE-downregulated genes have also been reported such as DUSP4^73,115^, which would be interesting to compare against our findings. To accomplish this (and also address our map’s ‘blind spot’ in the PHD1 domain) one might generate a variant effect map using single-cell RNA sequencing (scRNA-seq)^116^. This approach would benefit in that variant impacts on many different target genes could be assessed in parallel, rather than using insulin alone, and include more genes in closed chromatin. This approach would be limited by the stochastic nature of AIRE, such that different cells upregulate different genes^112^, and compounded by the limited number of independent reads that can be assayed in each cell using scRNA-seq. However, these limitations could be overcome by aggregating the results from ‘like’ sets of genes (e.g., genes for which there is *a priori* evidence for the assayed cell line that the promoter lies within closed chromatin).

Our APS-1 cohort-related analyses did not consider age, which is a limitation because APS-1 patients may develop more severe symptoms over time. Another limitation is that, given that *AIRE* is GC-rich and has reduced sequence coverage relative to other genes, we may not have detected all *AIRE* variants that are present for the *AIRE* patient cohort or UK BioBank participants.

Perhaps the most important use of our *AIRE* variant effect map will be as proactive (and therefore immediately available) functional evidence for classifying *AIRE* variants in suspected APS-1 patients, even for variants that have never before been clinically observed. This evidence promises to improve patient prognosis through earlier identification of patients who are at risk for APS-1 and may therefore benefit from earlier therapeutic intervention. Detection of pathogenic variants can also enable ‘cascade screening’ to identify first-degree relatives with the same variant who may also be at risk. Additionally, we have identified novel features of AIRE function that can lead to further studies and characterization, such as the impact of the first-reported variants that can increase reporter expression, and positions with deleterious impact that had not previously been reported. We have also identified a genotype-phenotype correlation in two patient cohorts. Thus, we provide the first *AIRE* variant effect map, contributing to *AIRE* clinical variant interpretation, to the knowledge of sequence-structure-function relationships, and also to the growing atlas of experimentally measured human variant effects^117^.

### Data and Code Availability

All files and scripts used for both analysis and figure generation are available at https://github.com/axakova/AIRE. AIRE functional scores, LLR_p_’s, evidence strengths and formal classifications are available in Data S2, and are also hosted on MaveDB^118^ (accession number: urn:mavedb:00001255-a-2).

## Acknowledgements

We gratefully acknowledge funding from the National Institute of Allergy and Infectious Diseases of the National Institutes of Health (R21AI173827), the National Institutes of Health National Human Genome Research Institute (NIH/NHGRI) Center of Excellence in Genomic Science Initiative (HG010461), the NIH/NHGRI Impact of Genomic Variation on Function (IGVF) Initiative (UM1HG011989), and a Canadian Institutes of Health Research Foundation Grant (FDN-159926) to F.P.R. We also acknowledge funding from the Novo Nordisk Foundation (NNF16OC0021254) and the Research Council of Norway (315599) to S.J.

## Supplemental Information

Document S1: Figures S1-S18, Tables S1-3, Supplemental Note, Supplemental Protocol Data S2: All datasets and primers

## References

1. Cinque, L. et al. Novel pathogenic variants of the AIRE gene in two Autoimmune Polyendocrine Syndrome Type I cases with atypical presentation: role of the NGS in diagnostic pathway and review of the literature. Biomedicines 8, (2020).

2. Husebye, E. S., Perheentupa, J., Rautemaa, R. & Kämpe, O. Clinical Manifestations and Management of Patients with Autoimmune Polyendocrine Syndrome Type I. J. Intern. Med. 265, 514–529 (2009).

3. Bjørklund, G., Pivin, M., Hangan, T., Yurkovskaya, O. & Pivina, L. Autoimmune polyendocrine syndrome type 1: Clinical manifestations, pathogenetic features, and management approach. Autoimmun. Rev. 21, 103135 (2022).

4. Landrum, M. J. et al. ClinVar: public archive of relationships among sequence variation and human phenotype. Nucl. Acids Res. 42, D980–D985 (2014).

5. Richards, S. et al. Standards and guidelines for the interpretation of sequence variants: a joint consensus recommendation of the American College of Medical Genetics and Genomics and the Association for Molecular Pathology. Genet. Med. 17, 405–424 (2015).

6. Starita, L. M. et al. Variant interpretation: functional assays to the rescue. Am. J. Hum. Genet. 101, 315–325 (2017).

7. Weile, J. et al. A framework for exhaustively mapping functional missense variants. Mol. Syst. Biol. 13, (2017).

8. Matreyek, K. A., Stephany, J. J. & Fowler, D. M. A platform for functional assessment of large variant libraries in mammalian cells. Nucleic Acids Res. 45, e102 (2017).

9. Gibson, D. G. et al. Enzymatic assembly of DNA molecules up to several hundred kilobases. Nat. Methods 6, 343–345 (2009).

10. Kean, M. J. et al. Structure-function analysis of core STRIPAK Proteins: a signaling complex implicated in Golgi polarization. J. Biol. Chem. 286, 25065–25075 (2011).

11. Tabet, D. R. et al. The functional landscape of coding variation in the familial hypercholesterolemia gene *LDLR*. Science (2025) doi:10.1126/science.ady7186.

12. Sun, S. et al. A proactive genotype-to-patient-phenotype map for cystathionine beta-synthase. Genome Med. 12, 13 (2020).

13. van Loggerenberg, W. et al. Systematically testing human HMBS missense variants to reveal mechanism and pathogenic variation. Am. J. Hum. Genet. 110, 1769–1786 (2023).

14. Axakova, A. et al. Landscapes of missense variant impact for human superoxide dismutase 1. The American Journal of Human Genetics 0, (2025).

15. Abramson, J. et al. Accurate structure prediction of biomolecular interactions with AlphaFold 3. Nature (2024) doi:10.1038/s41586-024-07487-w.

16. Chignola, F. et al. The solution structure of the first PHD finger of autoimmune regulator in complex with non-modified histone H3 tail reveals the antagonistic role of H3R2 methylation. Nucleic Acids Res. 37, 2951–2961 (2009).

17. Mitternacht, S. FreeSASA: An open source C library for solvent accessible surface area calculations. F1000Res. 5, 189 (2016).

18. Vangone, A. & Bonvin, A. M. Contacts-based prediction of binding affinity in protein-protein complexes. Elife 4, e07454 (2015).

19. Xue, L. C., Rodrigues, J. P., Kastritis, P. L., Bonvin, A. M. & Vangone, A. PRODIGY: a web server for predicting the binding affinity of protein-protein complexes. Bioinformatics 32, 3676–3678 (2016).

20. Weile, J. et al. Shifting landscapes of human MTHFR missense-variant effects. Am. J. Hum. Genet. 108, 1283–1300 (2021).

21. Wu, Y., Li, R., Sun, S., Weile, J. & Roth, F. P. Improved pathogenicity prediction for rare human missense variants. Am. J. Hum. Genet. 108, 1891–1906 (2021).

22. Epanechnikov, V. A. Non-parametric estimation of a multivariate probability density. Theory Probab. Appl. 14, 153–158 (1969).

23. Scott, D. W. On optimal and data-based histograms. Biometrika 66, 605–610 (1979).

24. Tavtigian, S. V. et al. Modeling the ACMG/AMP variant classification guidelines as a Bayesian classification framework. Genet. Med. 20, 1054–1060 (2018).

25. Tavtigian, S. V., Harrison, S. M., Boucher, K. M. & Biesecker, L. G. Fitting a naturally scaled point system to the ACMG/AMP variant classification guidelines. Hum. Mutat. 41, 1734–1737 (2020).

26. Bruserud, Ø. et al. A longitudinal follow-up of autoimmune polyendocrine syndrome type 1. J. Clin. Endocrinol. Metab. 101, 2975–2983 (2016).

27. Zaidi, G. et al. Autoimmune polyendocrine syndrome type 1 in an Indian cohort: a longitudinal study. Endocr. Connect. 6, 289–296 (2017).

28. Orlova, E. M. et al. Expanding the phenotypic and genotypic landscape of autoimmune polyendocrine syndrome type 1. J. Clin. Endocrinol. Metab. 102, 3546–3556 (2017).

29. Sato, K. et al. A novel missense mutation of AIRE gene in a patient with autoimmune polyendocrinopathy, candidiasis and ectodermal dystrophy (APECED), accompanied with progressive muscular atrophy: case report and review of the literature in Japan. Endocr. J. 49, 625–633 (2002).

30. Celmeli, F. et al. Unexplained cyanosis caused by hepatopulmonary syndrome in a girl with APECED syndrome. J. Pediatr. Endocrinol. Metab. 30, 365–369 (2017).

31. Pearce, S. H. et al. A common and recurrent 13-bp deletion in the autoimmune regulator gene in British kindreds with autoimmune polyendocrinopathy type 1. Am. J. Hum. Genet. 63, 1675–1684 (1998).

32. Yan, Z. et al. A case report and literature review: Identification of a novel AIRE gene mutation associated with Autoimmune Polyendocrine Syndrome Type 1 in East Asians. Medicine (Baltimore) 99, e20000 (2020).

33. Couturier, A., Saugier-Veber, P., Carel, J.-C., Bertherat, J. & Brézin, A. P. Keratopathy in autoimmune polyendocrinopathy syndrome type 1. Cornea 34, 1086–1091 (2015).

34. Mazza, C. et al. Clinical heterogeneity and diagnostic delay of autoimmune polyendocrinopathy-candidiasis-ectodermal dystrophy syndrome. Clin. Immunol. 139, 6–11 (2011).

35. Podkrajsek, K. T. et al. Detection of a complete autoimmune regulator gene deletion and two additional novel mutations in a cohort of patients with atypical phenotypic variants of autoimmune polyglandular syndrome type 1. Eur. J. Endocrinol. 159, 633–639 (2008).

36. Puzenat, E. et al. Diagnostic précoce du syndrome APECED : un défi pour le dermatologue. Ann. Dermatol. Venereol. 141, 290–294 (2014).

37. Saha, A., Kapadia, S. F., Vala, K. B. & Patel, H. V. Clinical utility of genetic testing in Indian children with kidney diseases. BMC Nephrol. 24, 212 (2023).

38. Sahoo, S. K. et al. Identification of autoimmune polyendocrine syndrome type 1 in patients with isolated hypoparathyroidism. Clin. Endocrinol. (Oxf.) 85, 544–550 (2016).

39. Meloni, A. et al. Two novel mutations of the AIRE protein affecting its homodimerization properties. Hum. Mutat. 25, 319 (2005).

40. Sanford, E., et al. Rapid whole-genome sequencing identifies a novel AIRE variant associated with autoimmune polyendocrine syndrome type 1. Cold Spring Harb Mol Case Stud 4, (2018).

41. Wang, Y. et al. Genetic screening in a large Chinese cohort of childhood onset hypoparathyroidism by next-generation sequencing combined with TBX1-MLPA. J. Bone Miner. Res. 34, 2254–2263 (2019).

42. F Magitta, N., et al. Autoimmune polyendocrine syndrome type I in Slovakia: relevance of screening patients with autoimmune Addison’s disease. Eur. J. Endocrinol. 158, 705–709 (2008).

43. Ward, L. et al. Severe autoimmune polyendocrinopathy-candidiasis-ectodermal dystrophy in an adolescent girl with a novel AIRE mutation: response to immunosuppressive therapy. J. Clin. Endocrinol. Metab. 84, 844–852 (1999).

44. Dominguez, M. et al. Autoimmune polyendocrinopathy-candidiasis-ectodermal dystrophy (APECED) in the Irish population. J. Pediatr. Endocrinol. Metab. 19, 1343–1352 (2006).

45. Pellegrino, M. et al. A novel homozygous mutation of the AIRE gene in an APECED patient from Pakistan: Case report and review of the literature. Front. Immunol. 9, 1835 (2018).

46. Cetani, F. et al. A novel mutation of the autoimmune regulator gene in an Italian kindred with autoimmune polyendocrinopathy-candidiasis-ectodermal dystrophy, acting in a dominant fashion and strongly cosegregating with hypothyroid autoimmune thyroiditis. J. Clin. Endocrinol. Metab. 86, 4747–4752 (2001).

47. Jafarpour, S. et al. Association of rare variants in genes of immune regulation with pediatric autoimmune CNS diseases. J. Neurol. 269, 6512–6529 (2022).

48. Abbott, J. K. et al. Dominant-negative loss of function arises from a second, more frequent variant within the SAND domain of autoimmune regulator (AIRE). J. Autoimmun. 88, 114–120 (2018).

49. Bellacchio, E. et al. The possible implication of the S250C variant of the autoimmune regulator protein in a patient with autoimmunity and immunodeficiency: in silico analysis suggests a molecular pathogenic mechanism for the variant. Gene 549, 286–294 (2014).

50. Oftedal, B. E. et al. Radioimmunoassay for autoantibodies against interferon omega; its use in the diagnosis of autoimmune polyendocrine syndrome type I. Clin. Immunol. 129, 163–169 (2008).

51. Stolarski, B. et al. Molecular background of polyendocrinopathy-candidiasis-ectodermal dystrophy syndrome in a Polish population: novel AIRE mutations and an estimate of disease prevalence. Clin. Genet. 70, 348–354 (2006).

52. Oftedal, B. E. et al. Dominant Mutations in the Autoimmune Regulator AIRE Are Associated with Common Organ-Specific Autoimmune Diseases. Immunity 42, 1185–1196 (2015).

53. Vogel, A. et al. Autoimmune regulator AIRE: evidence for genetic differences between autoimmune hepatitis and hepatitis as part of the autoimmune polyglandular syndrome type 1. Hepatology 33, 1047–1052 (2001).

54. Özer, Y. et al. Left ventricular systolic dysfunction related to adrenal insufficiency in a case due to autoimmune polyendocrine syndrome type 1 with a novel variant. Mol. Syndromol. 14, 65–70 (2023).

55. Brnich, S. E. et al. Recommendations for application of the functional evidence PS3/BS3 criterion using the ACMG/AMP sequence variant interpretation framework. Genome Med. 12, 3 (2019).

56. Ioannidis, N. M. et al. REVEL: An Ensemble method for predicting the pathogenicity of rare missense variants. Am. J. Hum. Genet. 99, 877–885 (2016).

57. Cheng, J. et al. Accurate proteome-wide missense variant effect prediction with AlphaMissense. Science 381, eadg7492 (2023).

58. Ilmarinen, T. et al. Functional interaction of AIRE with PIAS1 in transcriptional regulation. Mol. Immunol. 45, 1847–1862 (2008).

59. Meloni, A. et al. DAXX is a new AIRE-interacting protein. J. Biol. Chem. 285, 13012–13021 (2010).

60. Cai, C. Q., Zhang, T., Breslin, M. B., Giraud, M. & Lan, M. S. Both polymorphic variable number of tandem repeats and autoimmune regulator modulate differential expression of insulin in human thymic epithelial cells. Diabetes 60, 336–344 (2011).

61. Žumer, K., Low, A. K., Jiang, H., Saksela, K. & Peterlin, B. M. Unmodified Histone H3K4 and DNA-Dependent Protein Kinase Recruit Autoimmune Regulator to Target Genes. Mol. Cell. Biol. 32, 1354–1362 (2012-4).

62. Jin, P., Zhang, Q., Dong, C.-S., Zhao, S.-L. & Mo, Z.-H. A novel mutation in autoimmune regulator gene causes autoimmune polyendocrinopathy-candidiasis-ectodermal dystrophy. J. Endocrinol. Invest. 37, 941–948 (2014).

63. Sparks, A. E., Chen, C., Breslin, M. B. & Lan, M. S. Functional Domains of Autoimmune Regulator (AIRE) Modulate INS-VNTR Transcription in Human Thymic Epithelial Cells. J. Biol. Chem. 291, 11313–11322 (2016).

64. Org, T. et al. The Autoimmune Regulator PHD finger binds to non-methylated histone H3K4 to activate gene expression. EMBO Rep. 9, 370–376 (2008).

65. Palmer, J. P. et al. Insulin antibodies in insulin-dependent diabetics before insulin treatment. Science 222, 1337–1339 (1983).

66. Kuglin, B., Gries, F. A. & Kolb, H. Evidence of IgG autoantibodies against human proinsulin in patients with IDDM before insulin treatment. Diabetes 37, 130–132 (1988).

67. Giannopoulou, E. Z. et al. Islet autoantibody phenotypes and incidence in children at increased risk for type 1 diabetes. Diabetologia 58, 2317–2323 (2015).

68. Bansal, K., Yoshida, H., Benoist, C. & Mathis, D. The transcriptional regulator Aire binds to and activates super-enhancers. Nat. Immunol. 18, 263–273 (2017).

69. Abramson, J., Giraud, M., Benoist, C. & Mathis, D. Aire’s Partners in the Molecular Control of Immunological Tolerance. Cell 140, 123–135 (2010).

70. Yoshida, H. et al. Brd4 bridges the transcriptional regulators, Aire and P-TEFb, to promote elongation of peripheral-tissue antigen transcripts in thymic stromal cells. Proc. Natl. Acad. Sci. U. S. A. 112, E4448–E4457 (2015).

71. Shao, W., Zumer, K., Fujinaga, K. & Peterlin, B. M. FBXO3 Protein Promotes Ubiquitylation and Transcriptional Activity of AIRE (Autoimmune Regulator)*. J. Biol. Chem. 291, 17953–17963 (2016).

72. Huoh, Y.-S. et al. Dual functions of Aire CARD multimerization in the transcriptional regulation of T cell tolerance. Nat. Commun. 11, 1625 (2020).

73. Org, T. et al. AIRE activated tissue specific genes have histone modifications associated with inactive chromatin. Hum. Mol. Genet. 18, 4699–4710 (2009).

74. Matreyek, K. A. et al. Multiplex assessment of protein variant abundance by massively parallel sequencing. Nat. Genet. 50, 874–882 (2018).

75. Matreyek, K. A., Stephany, J. J., Chiasson, M. A., Hasle, N. & Fowler, D. M. An improved platform for functional assessment of large protein libraries in mammalian cells. Nucleic Acids Res. 48, e1–e1 (2020).

76. Kuroda, A. et al. Insulin Gene Expression Is Regulated by DNA Methylation. PLoS One 4, e6953 (2009).

77. Guo, H. H., Choe, J. & Loeb, L. A. Protein tolerance to random amino acid change. Proc. Natl. Acad. Sci. U. S. A. 101, 9205–9210 (2004).

78. Bowie, J. U., Reidhaar-Olson, J. F., Lim, W. A. & Sauer, R. T. Deciphering the message in protein sequences: tolerance to amino acid substitutions. Science 247, 1306–1310 (1990).

79. Jumper, J. et al. Highly accurate protein structure prediction with AlphaFold. Nature 596, 583–589 (2021).

80. Morgan, A. A. & Rubenstein, E. Proline: the distribution, frequency, positioning, and common functional roles of proline and polyproline sequences in the human proteome. PLoS One 8, e53785 (2013).

81. Gray, V. E., Hause, R. J. & Fowler, D. M. Analysis of large-scale Mutagenesis data to assess the impact of single amino acid substitutions. Genetics 207, 53–61 (2017).

82. Dunham, A. S. & Beltrao, P. Exploring amino acid functions in a deep mutational landscape. Mol. Syst. Biol. 17, e10305 (2021).

83. Finnish-German APECED Consortium. An autoimmune disease, APECED, caused by mutations in a novel gene featuring two PHD-type zinc-finger domains. Nat. Genet. 17, 399–403 (1997).

84. Ilmarinen, T. et al. The monopartite nuclear localization signal of autoimmune regulator mediates its nuclear import and interaction with multiple importin alpha molecules. FEBS J. 273, 315–324 (2006).

85. Bottomley, M. J. et al. The SAND domain structure defines a novel DNA-binding fold in transcriptional regulation. Nat. Struct. Biol. 8, 626–633 (2001).

86. Fang, Y., Bansal, K., Mostafavi, S., Benoist, C. & Mathis, D. AIRE relies on Z-DNA to flag gene targets for thymic T cell tolerization. Nature 628, 400–407 (2024).

87. Koh, A. S. et al. Aire employs a histone-binding module to mediate immunological tolerance, linking chromatin regulation with organ-specific autoimmunity. Proc. Natl. Acad. Sci. U. S. A. 105, 15878–15883 (2008).

88. Santos, J. C. et al. The AIRE G228W mutation disturbs the interaction of AIRE with its partner molecule SIRT1. Front. Immunol. 13, 948419 (2022).

89. Björses, P. et al. Mutations in the AIRE Gene: Effects on Subcellular Location and Transactivation Function of the Autoimmune Polyendocrinopathy-Candidiasis–Ectodermal Dystrophy Protein. Am. J. Hum. Genet. 66, 378–392 (2000-2).

90. Halonen, M. et al. APECED-causing mutations in AIRE reveal the functional domains of the protein. Hum. Mutat. 23, 245–257 (2004).

91. Pitkänen, J. et al. Cooperative activation of transcription by autoimmune regulator AIRE and CBP. Biochem. Biophys. Res. Commun. 333, 944–953 (2005).

92. Meloni, A., Incani, F., Corda, D., Cao, A. & Rosatelli, M. C. Role of PHD fingers and COOH-terminal 30 amino acids in AIRE transactivation activity. Mol. Immunol. 45, 805–809 (2008).

93. Pitkänen, J., Vähämurto, P., Krohn, K. & Peterson, P. Subcellular localization of the autoimmune regulator protein. characterization of nuclear targeting and transcriptional activation domain. J. Biol. Chem. 276, 19597–19602 (2001).

94. Bottomley, M. J. et al. NMR structure of the first PHD finger of Autoimmune Regulator Protein (AIRE1): Insights into Autoimmune Polyendocrinopathy-Candidiasis-Ectodermal Dystrophy (APECED) disease. J. Biol. Chem. 280, 11505–11512 (2005).

95. Huoh, Y.-S. et al. Mechanism for controlled assembly of transcriptional condensates by Aire. Nat. Immunol. 25, 1580–1592 (2024).

96. Varadi, M. et al. AlphaFold Protein Structure Database: massively expanding the structural coverage of protein-sequence space with high-accuracy models. Nucleic Acids Res. 50, D439–D444 (2022).

97. Mateos, B. et al. The Ambivalent Role of Proline Residues in an Intrinsically Disordered Protein: From Disorder Promoters to Compaction Facilitators. J. Mol. Biol. 432, 3093–3111 (2020).

98. Young, L., Jernigan, R. L. & Covell, D. G. A role for surface hydrophobicity in protein-protein recognition. Protein Sci. 3, 717–729 (1994).

99. Ng, P. C. & Henikoff, S. SIFT: Predicting amino acid changes that affect protein function. Nucleic Acids Res. 31, 3812–3814 (2003).

100. Brandes, N., Goldman, G., Wang, C. H., Ye, C. J. & Ntranos, V. Genome-wide prediction of disease variant effects with a deep protein language model. Nat. Genet. 55, 1512–1522 (2023).

101. Laisk, T. et al. Genome-wide association study identifies five risk loci for pernicious anemia. Nat. Commun. 12, 3761 (2021).

102. Chiou, J. et al. Interpreting type 1 diabetes risk with genetics and single-cell epigenomics. Nature 594, 398–402 (2021).

103. Eriksson, D. et al. GWAS for autoimmune Addison’s disease identifies multiple risk loci and highlights AIRE in disease susceptibility. Nat. Commun. 12, 959 (2021).

104. Bez, P., Ceraudo, M., Vianello, F., Rattazzi, M. & Scarpa, R. Where AIRE we now? Where AIRE we going? Curr. Opin. Allergy Clin. Immunol. 24, 448–456 (2024).

105. Berger, A. H. et al. High-resolution transcriptional impact of AIRE: effects of pathogenic variants p.Arg257Ter, p.Cys311Tyr, and polygenic risk variant p.Arg471Cys. Front. Immunol. 16, 1572789 (2025).

106. Su, M. A. et al. Mechanisms of an autoimmunity syndrome in mice caused by a dominant mutation in Aire. J. Clin. Invest. 118, 1712–1726 (2008).

107. Goldfarb, Y. et al. Mechanistic dissection of dominant AIRE mutations in mouse models reveals AIRE autoregulation. J. Exp. Med. 218, (2021).

108. Ilmarinen, T. et al. Functional analysis of SAND mutations in AIRE supports dominant inheritance of the G228W mutation. Hum. Mutat. 26, 322–331 (2005).

109. Oftedal, B. E. et al. Dominant-negative heterozygous mutations in AIRE confer diverse autoimmune phenotypes. iScience 26, 106818 (2023).

110. Sudlow, C. et al. UK biobank: an open access resource for identifying the causes of a wide range of complex diseases of middle and old age. PLoS Med. 12, e1001779 (2015).

111. Shirts, B. H., Pritchard, C. C. & Walsh, T. Family-Specific Variants and the Limits of Human Genetics. Trends Mol. Med. 22, 925–934 (2016).

112. Meredith, M., Zemmour, D., Mathis, D. & Benoist, C. Aire controls gene expression in the thymic epithelium with ordered stochasticity. Nat. Immunol. 16, 942–949 (2015-9).

113. Villaseñor, J., Benoist, C. & Mathis, D. AIRE and APECED: molecular insights into an autoimmune disease. Immunol. Rev. 204, 156–164 (2005).

114. Hasle, N. et al. High-throughput, microscope-based sorting to dissect cellular heterogeneity. Mol. Syst. Biol. 16, e9442 (2020).

115. Lovewell, T. R. J., McDonagh, A. J. G., Messenger, A. G., Azzouz, M. & Tazi-Ahnini, R. Meta-Analysis of Autoimmune Regulator-Regulated Genes in Human and Murine Models: A Novel Human Model Provides Insights on the Role of Autoimmune Regulator in Regulating STAT1 and STAT1-Regulated Genes. Front. Immunol. 9, 1380 (2018).

116. Xu, H. et al. Single cell sequencing as a general variant interpretation assay. Genomics (2023).

117. Fowler, D. M. et al. An Atlas of Variant Effects to understand the genome at nucleotide resolution. Genome Biol. 24, 147 (2023).

118. Esposito, D. et al. MaveDB: an Open-Source Platform to Distribute and Interpret Data from Multiplexed Assays of Variant Effect. Genome Biol. 20, 223 (2019).

