## Supplemental Files for "Systematic and proactive evaluation of AIRE missense variant effects"

### Supplemental Figures

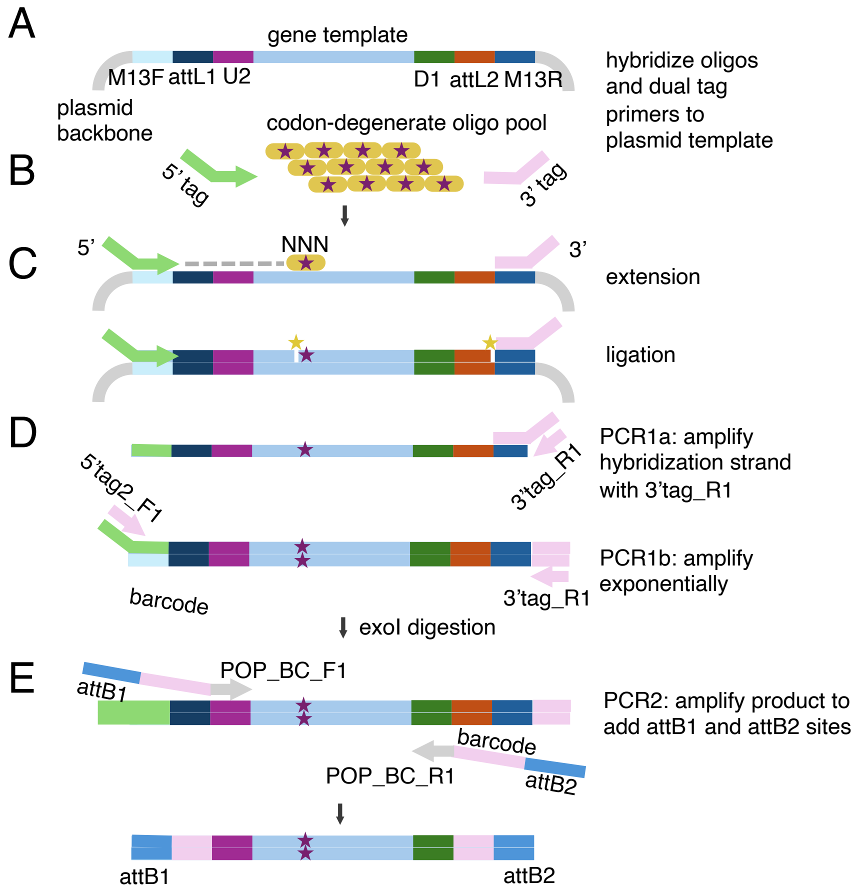


**Figure S1: Illustration of Dual-Tag POPCode method.**

A) The template used for Dual-Tag POPCode. The initial plasmid backbone contains M13F and M13R sites for sequencing, attL1 and attL2 sites remaining from Gateway cloning of the cDNA gene template into pDONR223, and universal U2 and D1 sites used for later addition of barcode and attB sites in PCR2.

B) To generate variants in the plasmid backbone, 5' and 3' tags and the oligo pool containing the 'NNN' degenerate codon are hybridized to the initial template in A.

C) Extension and ligation incorporate the oligo with the degenerate sequence to the template.

D) In PCR1, primers 5'tag2_F1 and 3'tag_R1 are used to preferentially amplify the mutagenized, hybridized strand, by only allowing amplification of the initial 3'tag. ExoI treatment rids the reaction of previous primers.

E) In PCR2, mutagenized products are amplified with POP_BC_F1 and POP_BC_R1 to add sequencing barcodes (if needed, depending on sequencing method) and attB1 and attB2 sites that are compatible with large-scale Gateway cloning.

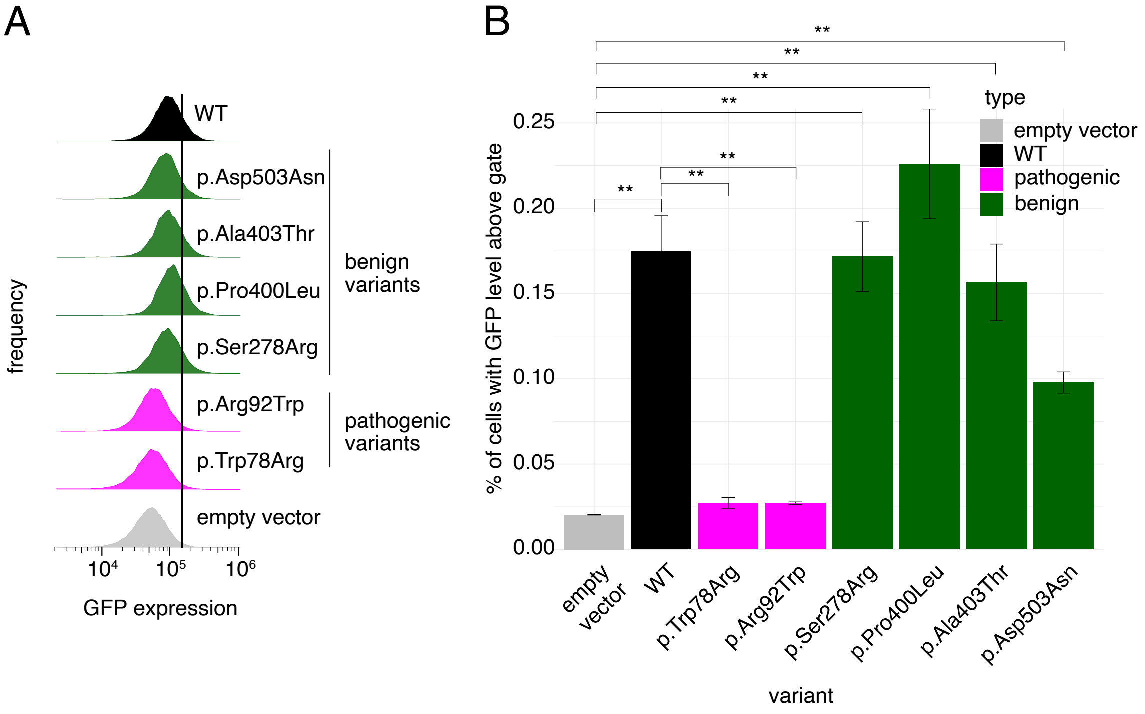


**Figure S2: Small-scale insulin promoter-GFP *AIRE* reporter assay results in HEK293 cells.**

A) GFP expression of known pathogenic or benign *AIRE* variants integrated into the *Bxb1* landing pad site of the HEK293 insulin promoter-GFP reporter cell line (Note: p.Ala403Thr has been reported as both benign and VUS). One representative replicate of three are shown.

B) Fraction of cells showing GFP expression from the HEK293 insulin promoter-GFP reporter assay that is above a gate threshold that was exceeded by 2% of cells in the empty vector control (mean of n=3 replicates). Error bars represent standard error, Welsh's *t*-test where "**" indicates $p<0.05$.

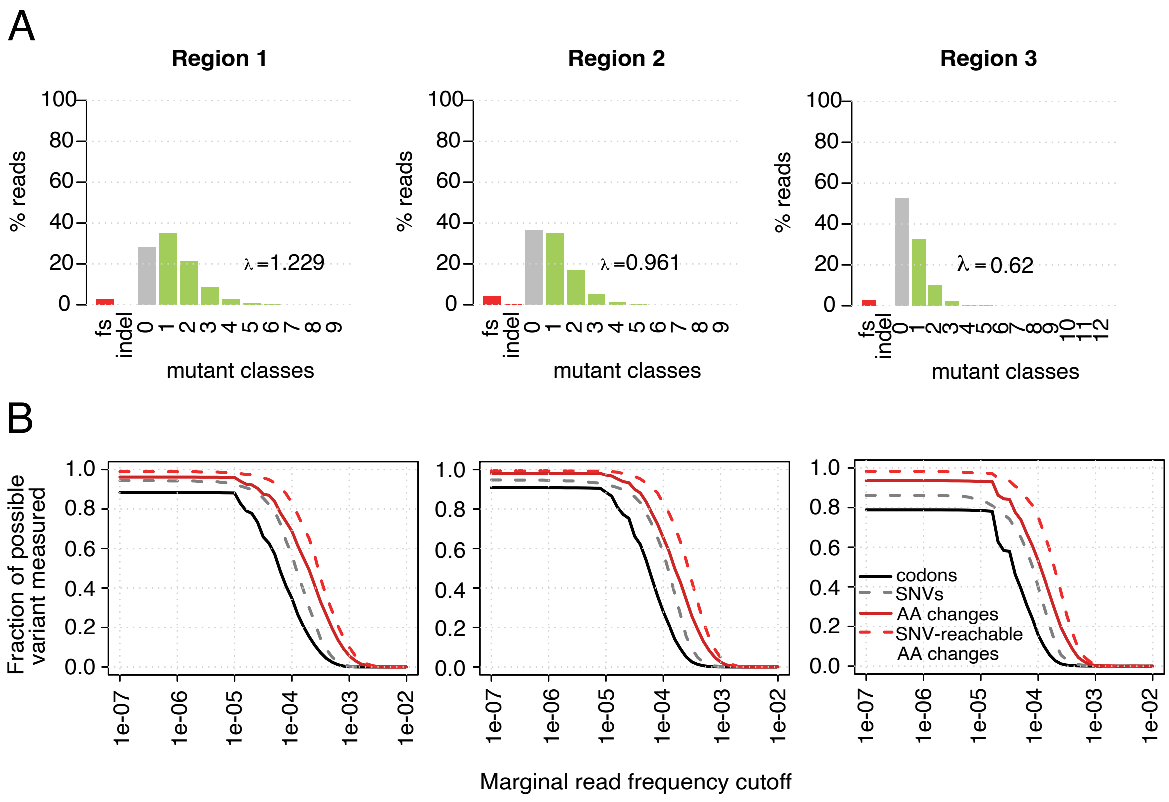


**Figure S3: *AIRE* plasmid pool variant library quality control.**

A) Representation of the number of variants/clone (λ) in the *AIRE* destination vector pool compatible with integration into the *Bxb1* landing pad site, by *AIRE* mutagenic region.

B) Amino acid coverage in the plasmid pool, including the fraction of: all possible codon-level changes (black line), single nucleotide-reachable amino acid changes (grey dotted line), all amino acid changes (red line), and SNV-reachable amino acid changes (red dotted line) based on read frequency.

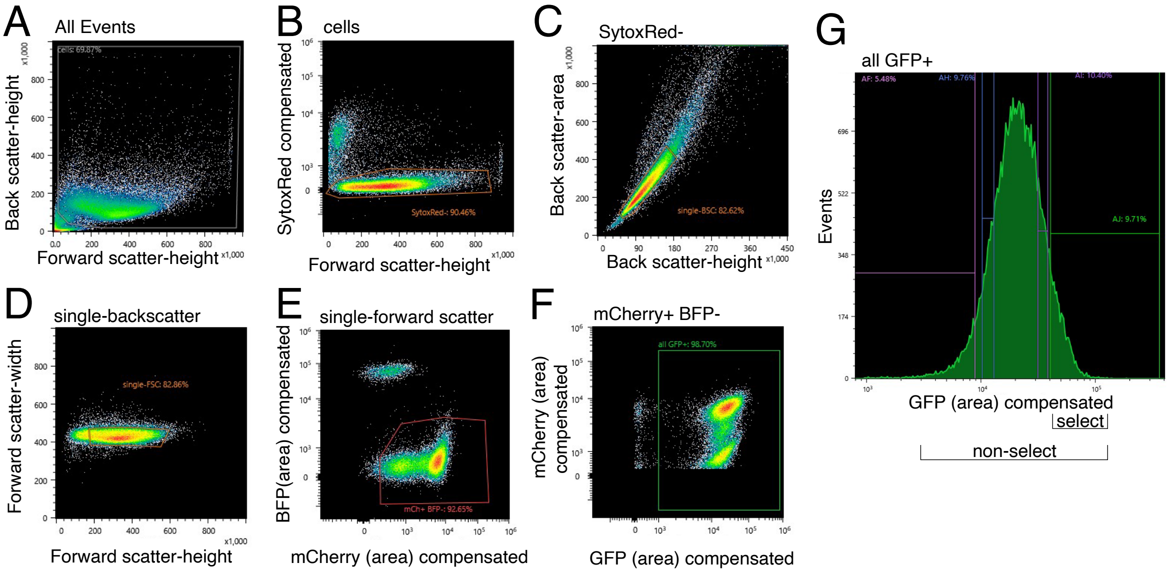


**Figure S4: Representative examples of cell sorting for cells expressing the insulin GFP reporter.**

A) Input cell population measuring back-scatter and forward-scatter for all sorting events, gated to exclude cellular debris at bottom left.

B) Cells from (A) gated for live cells measuring level of Sytox-red dead cell stain against forward scatter.

C) Live cells gated in (B) selected for singlets, using cell gating for back scatter height and back scatter area.

D) Additional single cell gating strategy of live single cells from (C) based on forward scatter width and height.

E) Selection for cells with *AIRE* integration events, using integration marker mCherry+ and lack of BFP signal from the *Bxb1* landing pad site.

F) Non-select cell sort, representative of all cells expressing the insulin promoter-GFP reporter.

G) Select cell sort, enriching for the cells which are found in the top 10% of insulin promoter-GFP expression. This sorting example is Region 3 of *AIRE*, but all three regions of *AIRE* had similar distributions.

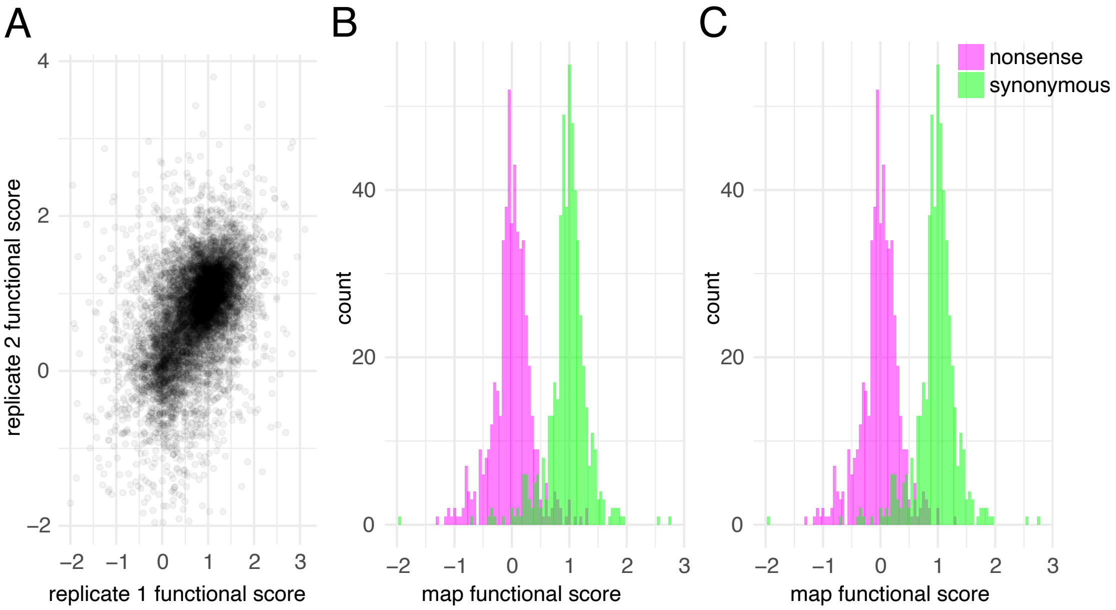


**Figure S5: Quality evaluation of *AIRE* variant effect map.**

A) Replicate agreement for *AIRE* variant scores between biological (transfection) replicates for insulin promoter-GFP reporter functional assay scores. Pearson's R=0.49, $p<2\times10^{-16}$.

B) The distribution of nonsense and synonymous *AIRE* variants, including nonsense variants at C-terminus of *AIRE*. Scores of synonymous variants were significantly higher than those of nonsense variants (Median scores of 1.00 and 0.01, respectively; $p<2\times10^{-16}$ by Wilcoxon test).

C) Distributions of nonsense and synonymous *AIRE* variants after excluding nonsense variants at the C-terminal 8 residue positions. (Median scores of synonymous and nonsense were 1.00 and 0.00, respectively; $p<2\times10^{-16}$ by Wilcoxon test).

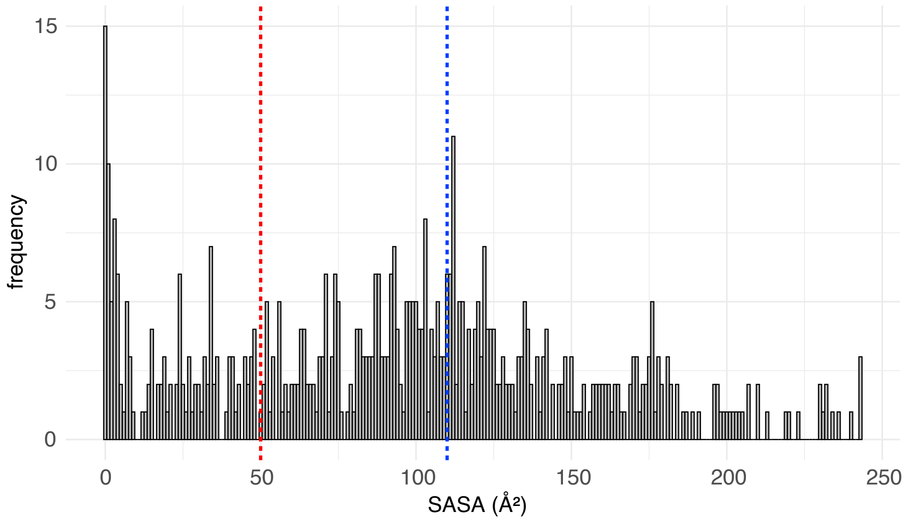


**Figure S6: Distribution of estimated surface accessibility for AIRE residues.**
Frequencies of estimated surface accessibility (calculated using FreeSASA scores[1]) are shown for residues in the reference AIRE protein. Red line indicates threshold below which positions are defined as buried, and blue line indicates threshold above which positions are defined as surface accessible. Here, ASA<50Å² defines buried and ASA>110Å² defines surface residues.

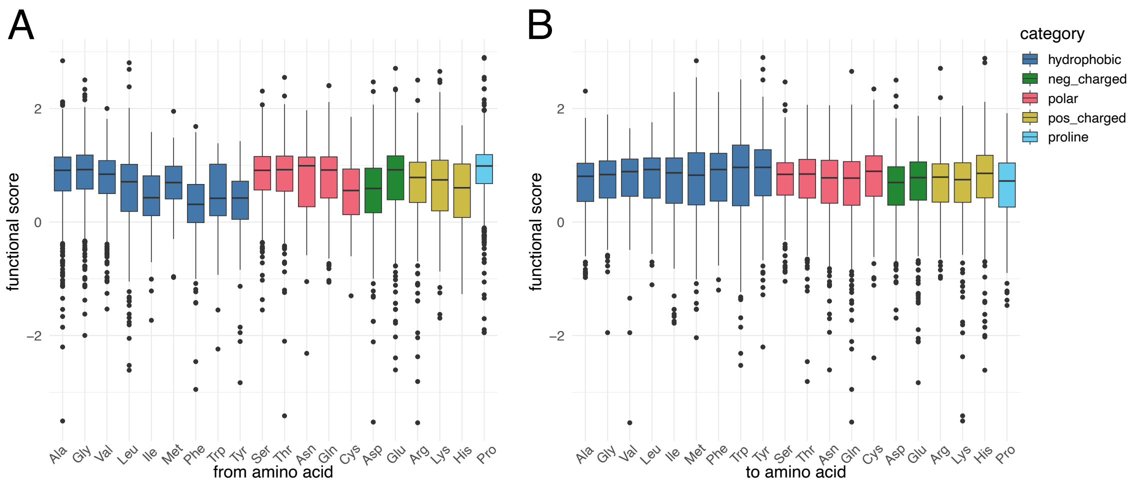


**Figure S7: Median functional scores of *AIRE* substitutions stratified by amino acid.**

A) Median functional scores for AIRE substitutions with the same reference (‘from’) amino acid. Boxes represent interquartile ranges. Whiskers either represent 1.5× the interquartile range or, in the absence of outlier points indicated by filled circles, they represent the minimum or maximum. Amino acids are colored by type (blue: hydrophobic, magenta: polar, green: negatively charged, yellow: positively charged, teal: proline.)

B) Median functional scores for AIRE substitutions with the same destination (‘to’) amino acid. Box and whisker plots are as described for (A).

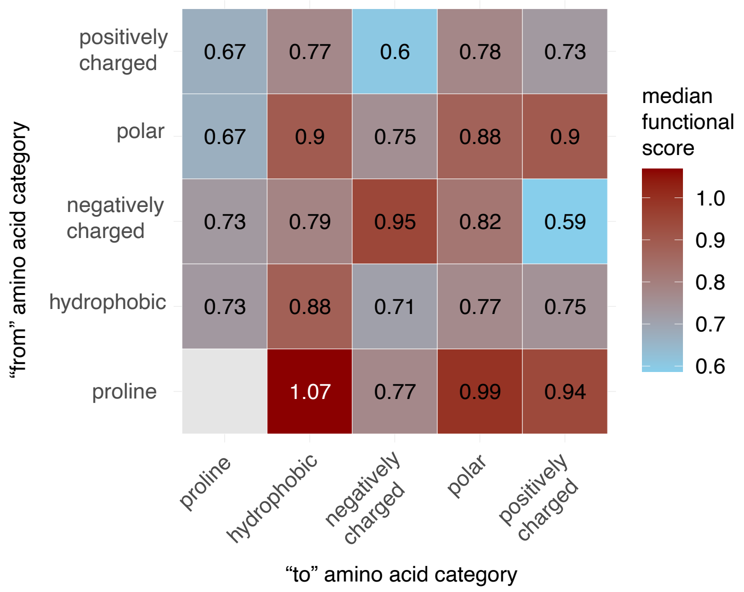


**Figure S8: Functional scores of missense substitutions grouped by reference ('from') and destination ('to') amino acid.** Amino acids were grouped into hydrophobic, negatively charged, positively charged, polar, and proline positions. Red and blue squares indicate higher (more tolerated) or lower (less tolerated) median functional scores, respectively. "Within-category" scores were included for all categories except proline (synonymous changes were excluded from these plots).

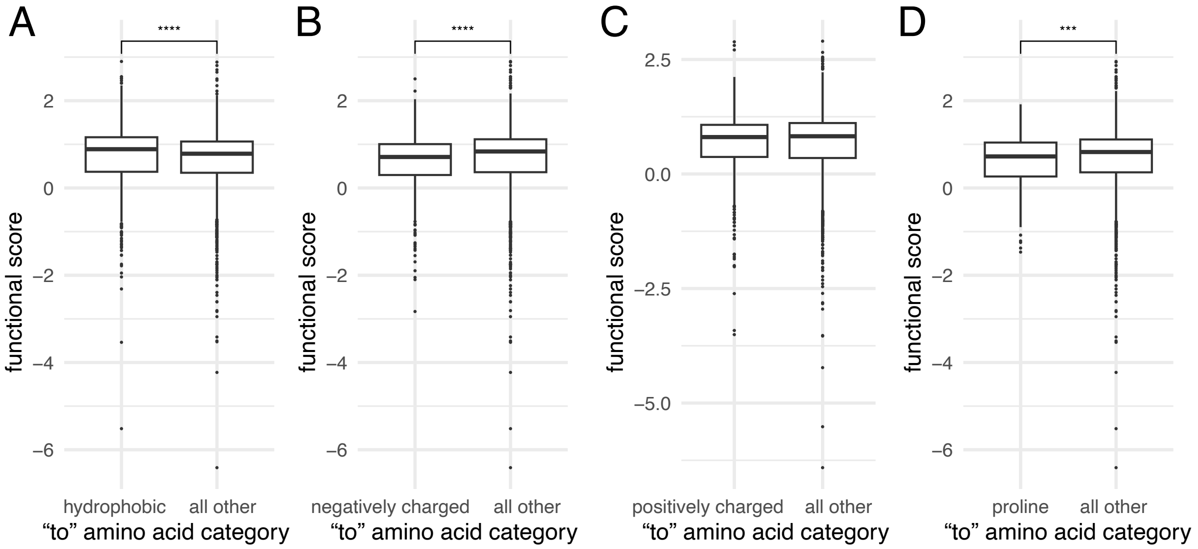


**Figure S9: Comparison of amino acid category-conserved substitution scores to all other non-conserved substitution scores.**

Median scores of substitutions within the same category as the reference ('from') amino acid, for categories: A) hydrophobic, B) negatively-charged, C) positively charged, and D) proline. Box and whisker plots are as described in Figure S7. Annotation with "***" indicates $p<1\times10^{-3}$, while "****" indicates $p<1\times10^{-4}$ by Wilcoxon test.

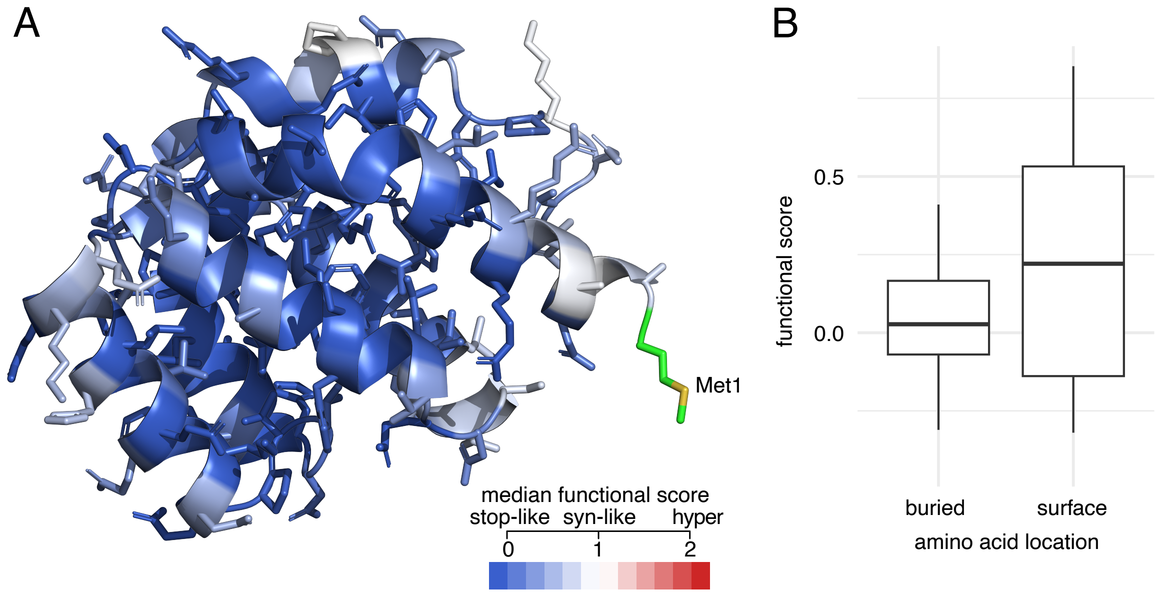


**Figure S10: Functional scores of the AIRE CARD domain.**

A) AIRE CARD domain "painted" by median score at each amino acid position onto AlphaFold 3 AIRE structure prediction. Blue indicates positions intolerant to variation, white indicates synonymous-like scores and red would indicate increased reporter signal if this had been observed for the CARD domain. The start codon is highlighted with Met1.

B) Scores of positions defined as buried or surface of the CARD domain according to FreeSASA scores (accessible surface area (ASA)<50Å² for buried, ASA>110Å² for surface). Here there was no significant difference between scores of buried and surface residues ($p>0.05$ by Wilcoxon test). Box and whisker plots are as described in Figure S7.

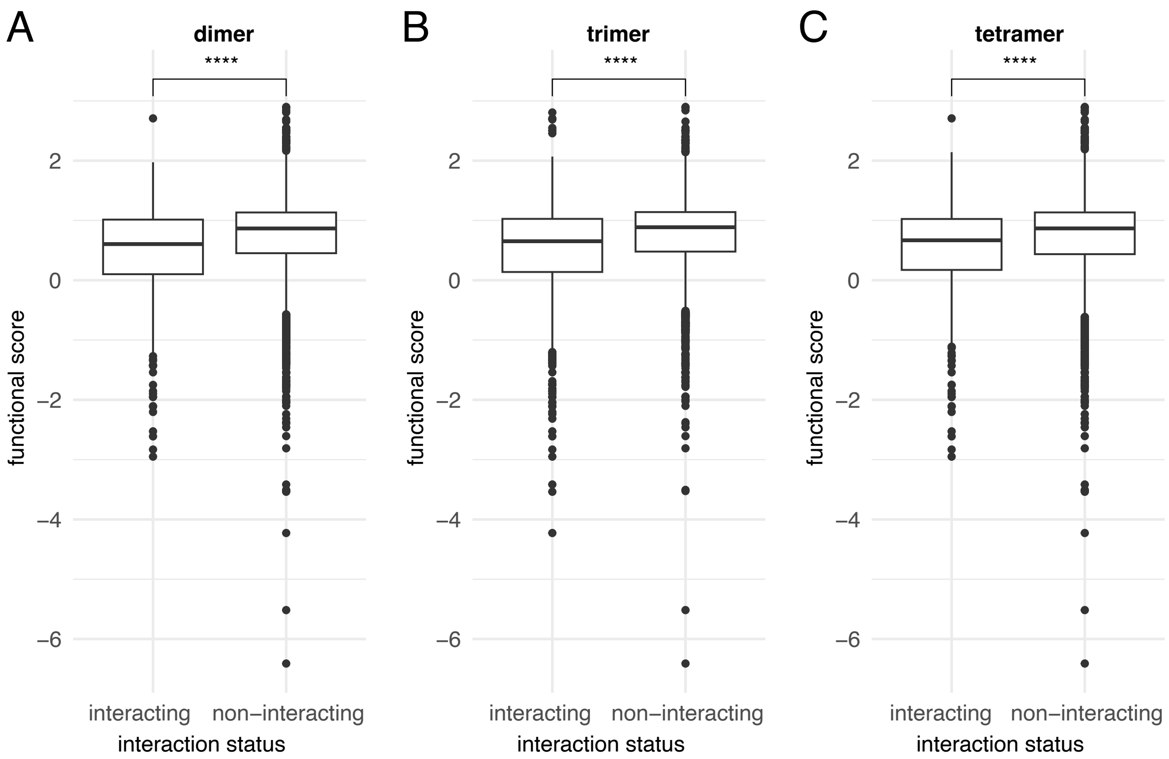


**Figure S11: Functional scores for interaction interfacial *AIRE* positions, identified via AlphaFold 3 co-fold models of dimerization, trimerization and tetramerization.**
Scores for positions identified to be interacting with other AIRE monomer(s) and those non-interacting as defined by Prodigy[2], for AIRE dimers (A), trimers (B) and tetramers (C) as modeled by AlphaFold 3[3]. Annotation of "****" indicates $p<1\times10^{-4}$ by Wilcoxon test.

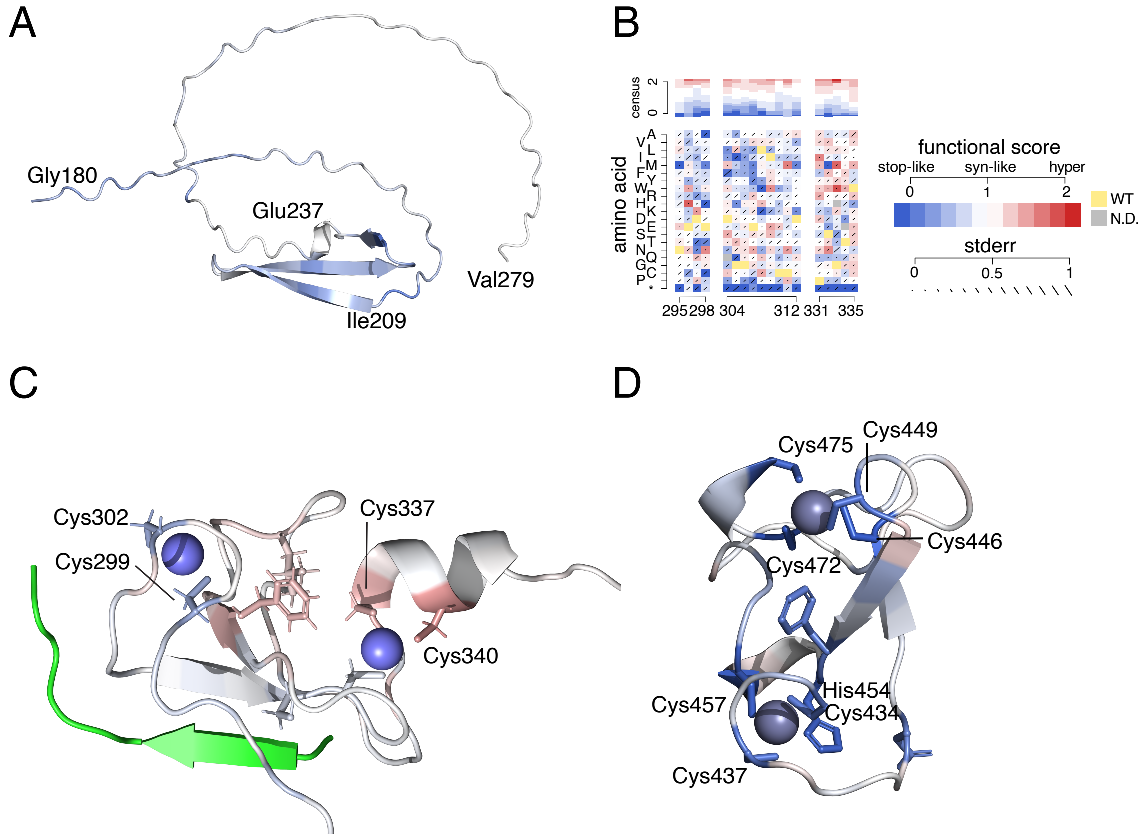


**Figure S12: Visualizing *AIRE* map scores in the context of predicted and experimental AIRE structure.**

A) Median map scores painted onto a predicted structure (AlphaFold 3) of AIRE's SAND domain.

B) *AIRE* variant effect map scores for known H3K4me0-interacting residues. WT residues are indicated in yellow, deleterious residues in blue, synonymous-like variants in white, and hyper (increased reporter expression) in red. Consensus track represents summary of positional effects.

C) Median *AIRE* variant effect scores were painted onto the PHD1 interaction with H3K4me0 NMR crystal structure PDB ID:2KE1[4]. H3K4me0 is painted in green. $Zn^{2+}$ ions are in blue, with $Zn^{2+}$-coordinating positions indicated.

D) Median variant effect map scores painted onto the structure (AlphaFold 3) of the AIRE PHD2 domain.

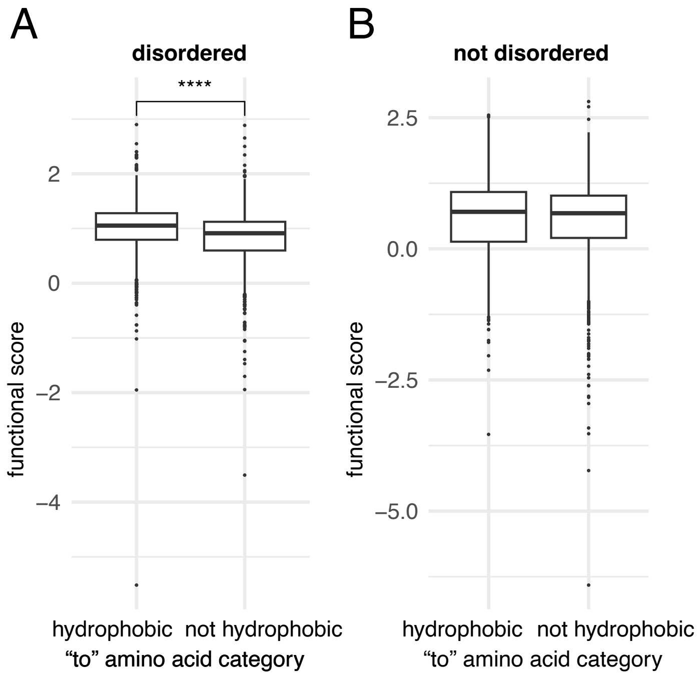


**Figure S13: AIRE variant effect trends related to hydrophobicity for disordered and non-disordered regions**

A) At AIRE positions known to be disordered, variant effects of changes to hydrophobic vs. non-hydrophobic residues, excluding hydrophobic to hydrophobic and non-hydrophobic to non-hydrophobic substitutions. "****" indicates $p<1\times10^{-4}$ by Wilcoxon test.

B) At AIRE positions not known to be disordered, variant effects of changes to hydrophobic vs. non-hydrophobic residues, excluding hydrophobic to hydrophobic and non-hydrophobic to non-hydrophobic substitutions. Here the difference was not significant ($p>0.05$ by Wilcoxon test).

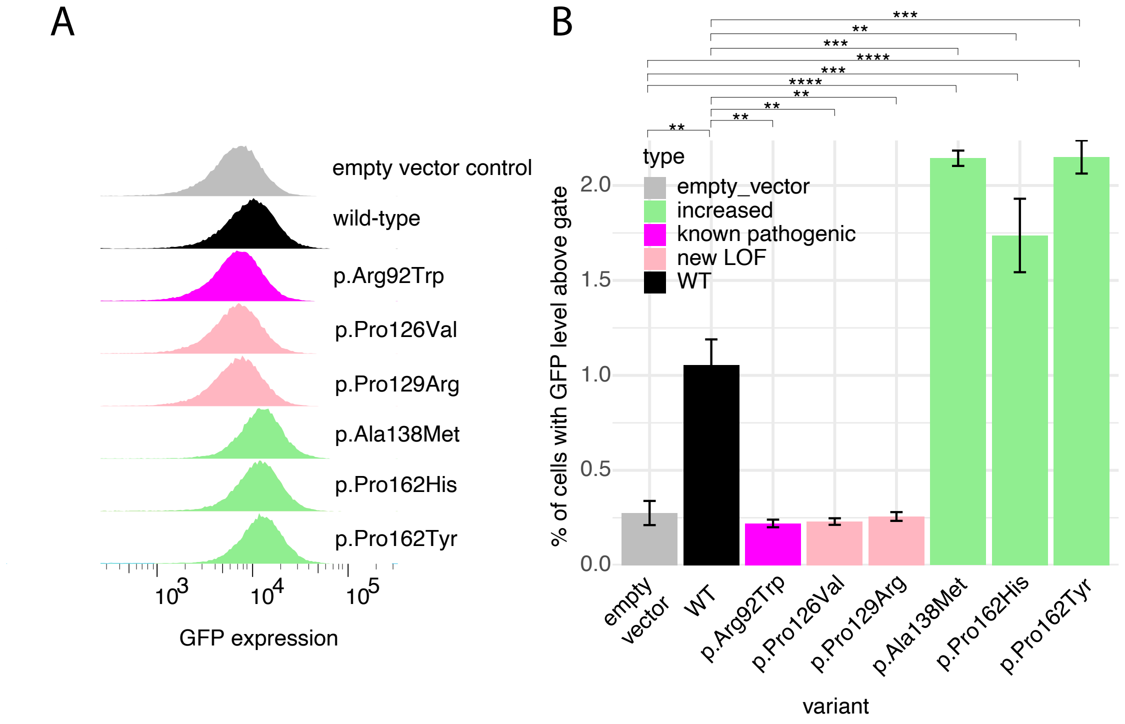


**Figure S14: Assessment of clonal *AIRE* variant impacts on insulin promoter-driven GFP expression.**

A) Representative flow cytometry plots of insulin promoter-GFP expression for HEK293 cell clones expressing individual *AIRE* variants (shown for one transfection replicate).

B) For the insulin promoter-GFP reporter assay, fraction of cells with GFP expression above a gate threshold set such that 1% of WT cells were above this threshold (n all variants = 3, n WT and empty vector = 2). Error bars represent standard error. "*", "**", "***", "****" indicates $p<0.05$, $5\times10^{-3}$, $5\times10^{-4}$, $5\times10^{-5}$, respectively, by Welsh's *t*-test.

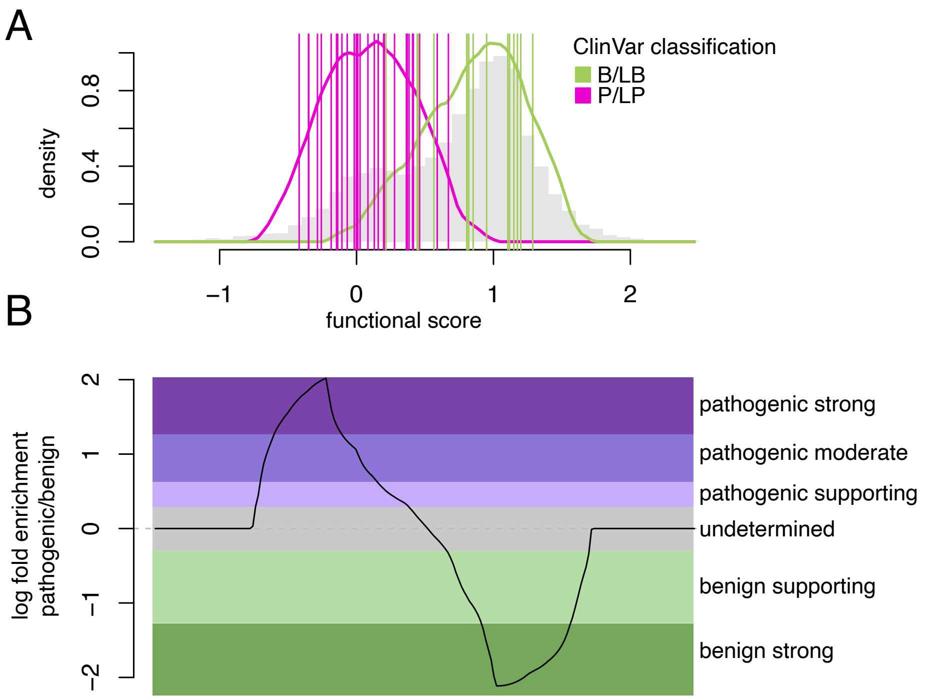


**Figure S15: Calibration to derive a log likelihood ratio of pathogenicity ($LLR_p$) corresponding to each *AIRE* variant effect functional score.**

A) Estimated score distributions of P/LP (magenta) and B/LB (green) variants. Vertical lines indicate individual scores for the 16 B/LB and 30 P/LP variants from ClinVar (excluding PHD1 reference set variants).

B) Calibrated $LLR_p$ and corresponding ACMG/AMP evidence strength labels for each *AIRE* missense variant effect map score.

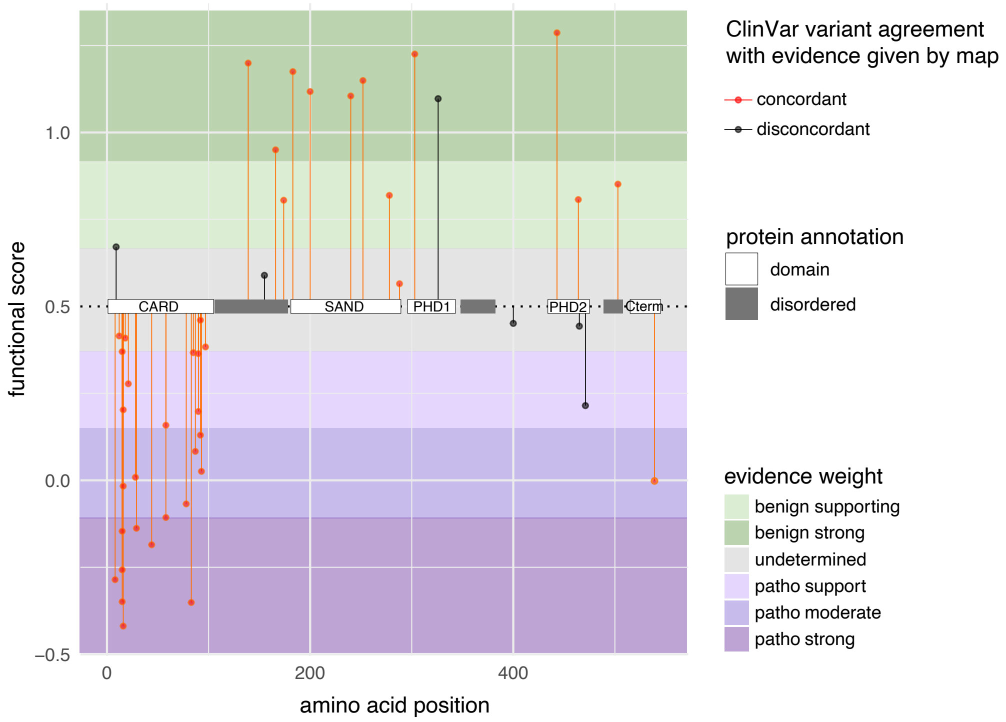


**Figure S16: Concordance of *AIRE* map score with pathogenicity annotations.**
Those map scores that are concordant with expectation (i.e. 'pointing in the right direction') are in red, while discordant scores are in black. Score thresholds indicating that ACMG/AMP evidence weight for classification should be provided are indicated by green towards benignity, purple towards pathogenicity, and grey for undetermined. AIRE protein domains are indicated by white boxes, with regions of disorder indicated with dark grey boxes.

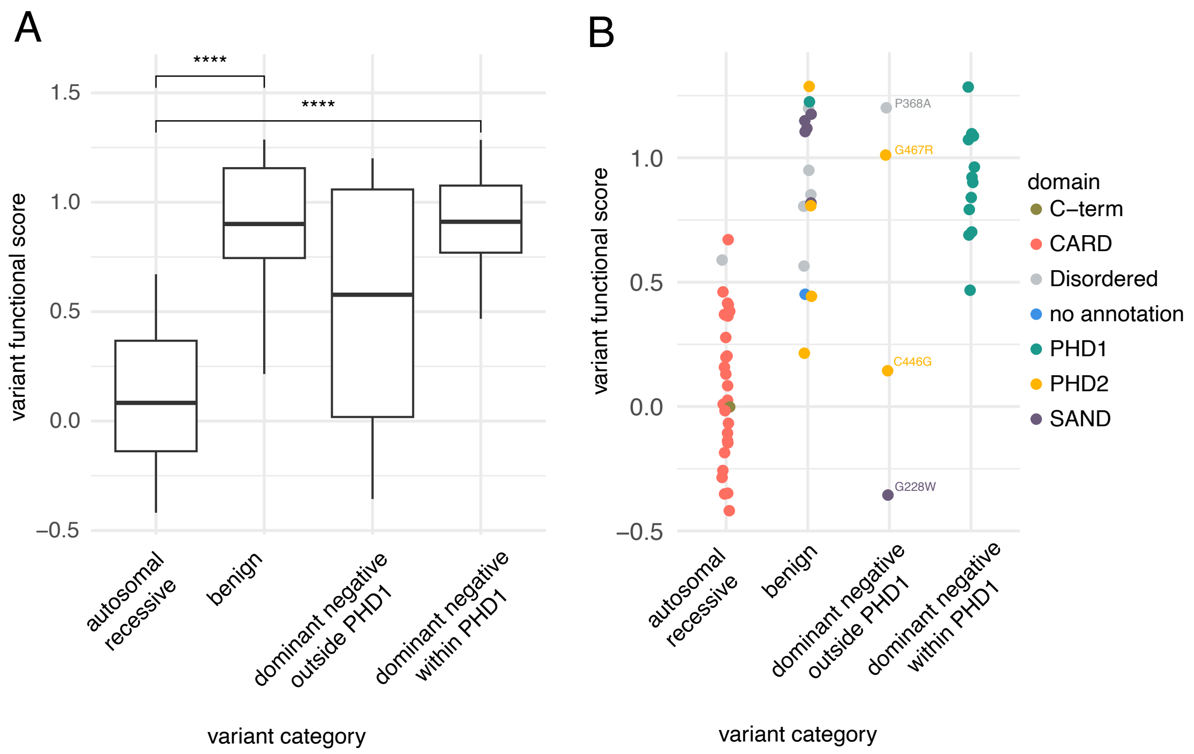


**Figure S17: *AIRE* variant effect map scores stratified by type of inheritance pattern.**

A) Scores of *AIRE* variants with known inheritance patterns. Dominant negative behavior is stratified based on whether or not the variant is within the PHD1 domain. Benign variants were included as a control. N autosomal recessive = 29, n benign = 16, n dominant negative outside PHD1 = 4, n dominant negative within PHD1 = 12. "****" indicates $p<1\times10^{-4}$ by Wilcoxon test.

B) Autosomal recessive and dominant negative variant map scores annotated by variant domain location and variants labelled if mentioned in text. Benign variants were included as a control.

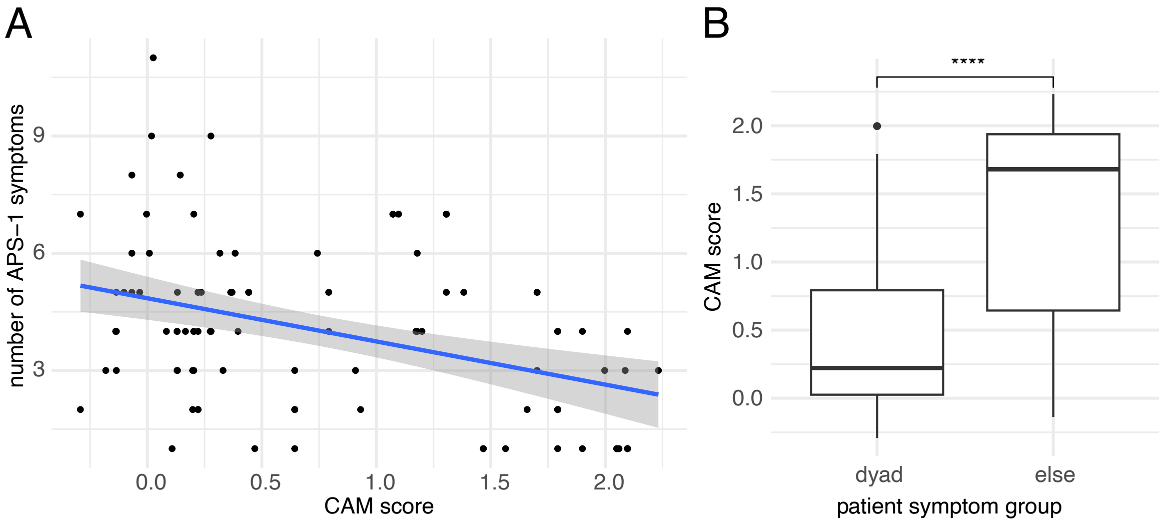


**Figure S18: Correspondence of combined-allele map scores with patient phenotypes.**

A) For an international cohort of APS-1 patients (n=97, including all patients with PHD1 variants), the relationship between *AIRE* combined-allele map (CAM) score (defined as the sum of functional scores for both patient alleles, where nonsense/frameshift and WT alleles are assigned scores of 0 and 1, respectively) and the number of APS-1 symptoms (R=-0.29, p=0.01).

B) Distributions of CAM scores for patients in the international APS-1 patient cohort who either do (n=61) or do not (n=36) exhibit the diagnostic dyad of APS-1 symptoms ($p=2\times10^{-7}$ by Wilcoxon test). Here we included patients with variants in the PHD1 domain.

**Supplemental Table 1: Variants which score significantly higher than the synonymous variant distribution, also indicating whether the amino acid substitution can be achieved via a single-nucleotide variant (i.e., is SNV accessible)**

| hgvs_pro | score | se | SNV accessible |
|---|---|---|---|
| p.Thr46Met | 2.55 | 0.11 | yes |
| p.Leu97Tyr | 2.69 | 0.14 | no |
| p.Pro109Met | 2.55 | 0.08 | no |
| p.Ala138Met | 2.84 | 0.08 | no |
| p.Pro162His | 2.88 | 0.06 | yes |
| p.Pro162Tyr | 2.90 | 0.10 | no |
| p.Pro315Trp | 2.51 | 0.08 | no |
| p.Lys395Met | 2.50 | 0.11 | yes |

**Supplemental Table 2: Variants with ClinVar annotations that disagree with the direction of evidence indicated by the map score.**

| hgvs_pro | score | se | Map-derived evidence weight | type | domain | SNV accessible |
|---|---|---|---|---|---|---|
| p.Arg9Pro | 0.67 | 0.54 | benign.support | P/LP | CARD | yes |
| p.Gly155Ser | 0.59 | 0.56 | undetermined | P/LP | N/A | yes |
| p.Pro326Leu | 1.10 | 0.25 | undetermined | P/LP | PHD1 | yes |
| p.Pro400Leu | 0.45 | 0.31 | undetermined | B/LB | N/A | yes |
| p.Arg465Gln | 0.44 | 0.20 | undetermined | B/LB | PHD2 | yes |
| p.Arg471Cys | 0.21 | 0.10 | patho.support | B/LB | PHD2 | yes |

**Supplemental Table 3: Participant symptom frequencies in International patient cohort.**

| Phenotype | count |
|---|---|
| Hypoparathyroidism | 65 |
| Chronic mucocutaneous candidiasis | 56 |
| Addison's disease/primary adrenal failure | 54 |
| Alopecia | 22 |
| Enamel hypoplasia | 22 |
| Hypothyroidism | 19 |
| Primary ovarian insufficiency | 19 |
| Pernicious Anemia | 18 |
| Type 1 diabetes | 14 |
| Malabsorption | 13 |
| Chronic active hepatitis | 11 |
| Vitiligo | 11 |
| Nail dystrophy | 10 |
| Keratitis or keratopathy | 7 |
| Chronic active gastritis | 5 |
| Epilepsy | 3 |
| Red cell aplasia | 3 |
| Asplenia or hyposplenia | 3 |
| Testicular failure | 3 |
| Cataract | 2 |
| Interstitial lung disease | 2 |
| Juvenile arthritis | 2 |
| Myopathy | 2 |
| Acute disseminated encephalomyelitis | 1 |
| Autoimmune encephalitis | 1 |
| Blepharitis | 1 |
| Chronic diarrhea | 1 |
| Viral encephalitis | 1 |
| Exocrine pancreatic failure | 1 |
| Fibroma auricular | 1 |
| Periodic fever with rash | 1 |
| Growth hormone deficiency | 1 |
| Growth retardation | 1 |
| Gingivitis | 1 |
| Chronic active hepatitis | 1 |
| Hearing loss | 1 |
| Hypophysitis | 1 |
| Hypogammaglobulinemia | 1 |
| Metaphyseal dysplasia | 1 |
| Myelin oligodendrocyte glycoprotein antibody-associated di | 1 |

| | |
|---|---|
| Chronic otitis media | 1 |
| Ptosis | 1 |
| Rheumatoid arthritis | 1 |
| Retinitis pigmentosa | 1 |
| Squamous cell carcinoma of the oral mucosa | 1 |
| Urticaria | 1 |

**Supplemental Note**
Insulin core promoter sequence:
5’-
gtggggacaggggtctggggacagcagcgcaaagagccccgccctgcagcctccagctctcctggtctaatgtggaaagtggcccaggtg
agggctttgctctcctggagacatttgcccccagctgtgagcagggacaggtctggccaccgggcccctggttaagactctaatgacccgctgg
tcctgaggaagaggtgctgacgaccaaggagatcttcccacagacccagcaccagggaaatggtccggaaattgcagcctcagcccccca
gccatctgccgacccccccaccccaggccctaatgggccaggcggcaggggttgagaggtaggggagatgggctctgagactataaagc
cagcgggggcccagcagccctcagccctccaggacaggctgcatcag-3’

*AIRE* codon-optimized coding sequence:
5’ATGGCGACGGACGCGGCGCTACGCCGGCTTCTGAGGCTCCATCGTACCGAAATTGCAGTTGC
AGTCGATTCCGCTTTTCCCCTTCTTCATGCTCTCGCAGATCATGATGTTGTACCGGAAGATAAATT
CCAAGAAACTTTGCACTTGAAAGAGAAAGAAGGGTGTCCACAAGCTTTTCATGCGCTGCTTAGCT
GGCTTCTTACACAAGATTCAACCGCAATACTTGATTTCTGGCGGGTCTTGTTTAAAGATTATAATC
TTGAACGGTACGGGCGCTTGCAACCTATTCTTGATAGTTTTCCTAAGGACGTCGATTTGTCTCAA
CCTAGAAAAGGCCGGAAACCGCCCGCAGTACCGAAAGCGCTGGTCCCACCCCCTCGCCTGCCG
ACTAAAAGAAAAGCTAGTGAGGAAGCAAGGGCCGCTGCTCCTGCCGCTTTGACCCCTCGAGGTA
CAGCTAGTCCTGGATCCCAGCTCAAAGCTAAACCGCCTAAGAAACCAGAATCTTCCGCCGAACA
ACAAAGACTGCCCCTGGGAAATGGCATACAAACAATGAGCGCCAGCGTTCAACGGGCCGTAGCT
ATGTCTTCTGGCGATGTTCCTGGTGCACGGGGAGCTGTAGAAGGAATTCTTATACAACAAGTCTT
CGAAAGTGGTGGGAGTAAGAAATGTATTCAAGTGGGCGGCGAATTTTATACACCTTCAAAATTTG
AGGATAGTGGATCTGGCAAGAATAAAGCTAGATCTTCCTCAGGTCCAAAACCACTTGTGAGAGC
GAAAGGTGCACAAGGGGCGGCGCCTGGCGGCGGCGAAGCACGCTTGGGTCAACAAGGATCCG
TGCCAGCTCCACTCGCTCTTCCGAGCGATCCGCAACTGCATCAAAAGAACGAAGATGAATGCGC
TGTATGCCGCGATGGCGGTGAACTGATTTGTTGCGATGGATGTCCCAGGGCTTTTCATTTGGCG
TGTCTCTCTCCGCCTTTGAGGGAAATACCATCTGGTACTTGGCGCTGTTCTTCCTGTCTCCAAGC
TACCGTGCAAGAAGTCCAACCACGCGCTGAAGAACCGAGACCACAAGAACCCCCTGTCGAAACA
CCCCTTCCACCTGGCTTGAGATCAGCCGGTGAAGAAGTGCGGGGCCCACCCGGCGAGCCGTTG
GCGGGAATGGATACTACCCTCGTGTATAAACATTTGCCAGCACCACCCAGCGCTGCGCCTTTGC
CGGGCCTCGATAGCTCAGCTTTGCATCCGCTTCTTTGCGTAGGCCCCGAAGGGCAACAAAATTT
GGCCCCCGGAGCACGCTGTGGAGTTTGTGGGGACGGAACCGATGTTCTTAGATGCACCCATTG
TGCAGCAGCTTTTCATTGGCGATGTCATTTTCCTGCGGGTACAAGCCGCCCTGGCACCGGGCTT
CGATGTCGGAGCTGTTCTGGCGATGTTACTCCTGCACCGGTCGAAGGAGTACTCGCTCCAAGTC
CTGCACGGTTGGCGCCCGGACCCGCTAAAGACGATACGGCTTCCCATGAACCAGCGCTCCATA
GAGACGATCTCGAATCACTGTTGAGTGAACATACATTTGACGGGATTCTCCAATGGGCTATACAA
AGTATGGCAAGGCCTGCCGCACCATTCCCCTCCTGA 3’.

**Supplemental Protocol**

**Dual-tag POPCode protocol (See Figure S1).**

**Purpose**: To generate a library of missense variants for a gene of interest, compatible for large-scale Gateway BP cloning into pDONR223.

**Notes and suggestions**:

- Do not stop until after PCR1 because products are single-stranded until then.
- Phosphorylation reactions can be done in advance but freshly prepared reactions may be more efficient. Use fresh ATP only.

**Primer list:**

| **Name** | **Sequence** | **Step** |
|---|---|---|
| M13ext_5'Tag2_F | GCATGCCAATACTGGTAGATCACCGTAAAACGAC GGCCAGTCTTAA | 2 |
| 3'Tag_Hyb | ACATGGTCATAGCTGTTTCCTGGCACGACCGCTCT ATTACTTAGAG | 2 |
| 5'tag2_F1 | GCATGCCAATACTGGTAGATCACC | 5 (PCR1) |
| 3'TagR_amp | CTCTAAGTAATAGAGCGGTCGTGC | 5 (PCR1) |
| POP_BC_F1 (PAGE purified) | GGGGACAACTTTGTACAAAAAAGCAGGCTCCATAC GAGCACATTACGGGSWSWSWSWSWSWSWSWS WSWSWSWSCTAACTCGCATACCTCTGATAAC | 7 (PCR2) |
| POP_BC_R1 (PAGE purified) | GGGGACAACTTTGTACAAGAAAGCTGGGTGCTTG ACTGAGCGACTGAGGSWSWSWSWSWSWSWSWS WSWSWSWSCCTTCACACGCACCTATCGAAGTCA | 7 (PCR2) |

**Materials list:**

1. Gene for mutagenesis with stop codon in pDONR223, including U2 and D1 constant regions for barcode addition in PCR2 (Figure S1A).
2. Mutagenic oligos with degeneracy centered on each codon, specific to the gene. These should be ordered in buffer to a concentration of 100uM for each.
3. PNK and PNK buffer (NEB).
4. 2X Phusion MM.
5. Taq DNA ligase buffer and Taq DNA ligase.
6. Exonuclease I (NEB).

**Step 1: Phosphorylate oligos and 3'Tag_Hyb**

*Purpose*: Mutagenic oligos need to be phosphorylated to provide a substrate for DNA ligase.

1. Pool 4uL of each mutagenic 100uM oligo per region.
2. Phosphorylate oligos and 3'Tag_Hyb (do many 3'tag ones - calculate how much is needed if you have a lot of regions/conditions. 5uL of phosphorylated 3'Tag_Hyb is needed per hybridization reaction).

1X phosphorylation reaction:

| Material: | Amount (uL) |
|---|---|
| 10X PNK buffer (NEB) | 5 |
| 10mM ATP | 5 |
| 100uM oligo pool (300 pmol) or 3'Tag_Hyb | 3 |
| PNK (NEB) | 1 |
| H2O | 36 |

3. Oligos from this reaction are 6uM final.
4. Prepare phosphorylation reaction, flick tube, quick spin.
5. Incubate 1 hour at 37C.

**Step 2: Hybridization (Figure S1B)**

*Purpose*: Hybridize mutagenic oligos to template.

The concentration of oligos needed for mutagenesis between 0.4-1 mutations per clone will depend on the target, and three concentrations should be tested to start. Usually, 3-6n is appropriate but may depend on the gene.

n = 2 x (number of unique oligos per region/150)

Example: Region 1 of AIRE consists of the first 182 amino acids of the protein (meaning 182 unique mutagenic oligos).

for AIRE: n= 2x(180)/150 = 2.4 uL

1. Combine the following in a PCR tube:

| **Material:** | **1X** |
|---|---|
| Gene you wish to mutagenize in pDONR223 with U2 and D1 regions | 90 ng |
| M13Fext_5'Tag2_F (10uM) | 3 |
| Phosphorylated 3'Tag_Hyb (6uM) | 5 |
| Phosphorylated oligos (6uM) | n |
| $H_2O$ | up to 20 |
| Total | 20 |

2. Flick and spin down tubes
3. Denature this reaction in thermal cycler at 95°C for 3 min then cool to 4°C for 15 min in thermocycler block. The oligos anneal as the temperature drops. If using a standard PCR block, the default ramp speed should be used. Lowering ramp speed was determined to be not as efficient.
4. Heat inactivate 2X Phusion Master Mix. Put 5uL phusion into a different thermocycler block 95°C for 3 min then cool to 4°C. Do not vortex after this step, flick and spin down.

**Step 3: Extension (Figure S1C)**

*Purpose*: Fill-in mutagenized strand.

1. Add 5ul of the oligo hybridization reaction to 5ul of heat activated 2X Phusion MM.
2. Incubate 2 hours at 50°C.

**Step 4: "Nick" ligation**

*Purpose*: Ligate the gaps in the amplified strand.

| **Material:** | **1x (uL)** |
|---|---|
| Taq DNA ligase buffer | 1.5 |
| Taq ligase | 0.5 |
| $H_2O$ | 3 |
| Total | 5 |

1. Scale this reaction to required volume, and add 5 uL to each amplification reaction.
2. Incubate 20min at 45°C. (Note: Ligation for 1 hour removed smaller-than-expected non-specific bands in subsequent steps for some genes).

**Step 5: PCR1 - Tag-based amplification (Figure S1D)**
*Purpose*: Amplify the mutagenized strand preferentially.

| **Material:** | **1X (uL)** |
|---|---|
| 1 uL ligated product template | 1 |
| 25uL 2x Phusion MM | 25 |
| 10uM 5'tag2_F1 | 2 |
| 3'TagR_amp | 2 |
| $H_2O$ | 20 |
| Total | |

Thermal cycler conditions for PCR1:

| **Step** | **Temperature** | **Time** |
|---|---|---|
| 1 | 98°C | 30sec |
| 20 cycles of steps 2-4: | | |
| 2 | 98°C | 30sec |
| 3 | 65°C | 30sec |
| 4 | 72°C | 2min30sec |
| 5 | 72°C | 5min |
| 6 | 10°C | Forever |

**Step 6: Prevent primer carry-over**
*Purpose*: This step is important to remove the previous Tag-based amplification oligos. Exonuclease I digestion was more effective than PCR purification columns to remove primers.
ExoI digestion:

1. Add 5uL of exonuclease I/50 uL PCR reaction. Alternatively, exonuclease digest 10 uL of product with 1 uL of exonuclease I.
2. Incubate at 37°C for 15 minutes.
3. Deactivate by incubating at 80°C for 15 minutes.

**Step 7: PCR2 - Barcode and attB addition (Figure S1E)**
*Purpose*: Amplify Tag-based product with primers that add barcodes and attB sites for cloning.

1. Pool 4-5X reactions for each oligo concentration condition.

| **Material** | **1X (uL)** |
|---|---|
| Exonuclease I-digested product | 1uL (approximately 28 ng) |
| Phusion | 25 |
| POP-BC_F1 (100uM stock directly) | 2 |
| POP-BC_R1 (100uM stock directly) | 2 |
| $H_2O$ | 20 |
| Total | |

2. Thermal cycler conditions for attB addition:

| **Step** | **Temperature** | **Time** |
|---|---|---|
| 1 | 98°C | 30sec |
| 5 cycles of steps 2-4: | | |
| 2 | 98°C | 15sec |
| 3 | 58°C | 30sec |
| 4 | 72°C | 2min30sec |
| 12 cycles of steps 5-6: | | |
| 5 | 98°C | 15sec |
| 6 | 72°C | 2min30sec |

| | | |
|---|---|---|
| 7 Extension | 72°C | 5min |
| 8 Hold | 10°C | forever |

3. Pool products of the same concentration together, run on a 1% gel with combined wells.
4. Gel extract product with normal Qiagen gel extraction kit.
5. Quantify library with Qubit.
6. Ready for tiling and indexing for library QC.
7. This library is compatible with large-scale BP reactions into pDONR223.

**Supplemental References**


1. Mitternacht, S. FreeSASA: An open source C library for solvent accessible surface area calculations. *F1000Res.* **5**, 189 (2016).

2. Xue, L. C., Rodrigues, J. P., Kastritis, P. L., Bonvin, A. M. & Vangone, A. PRODIGY: a web server for predicting the binding affinity of protein-protein complexes. *Bioinformatics* **32**, 3676–3678 (2016).

3. Abramson, J. *et al.* Accurate structure prediction of biomolecular interactions with AlphaFold 3. *Nature* (2024) doi:10.1038/s41586-024-07487-w.

4. Chignola, F. *et al.* The solution structure of the first PHD finger of autoimmune regulator in complex with non-modified histone H3 tail reveals the antagonistic role of H3R2 methylation. *Nucleic Acids Res.* **37**, 2951–2961 (2009).
